# Conserved lipid and small molecule modulation of COQ8 reveals regulation of the ancient UbiB family

**DOI:** 10.1101/149823

**Authors:** Andrew G. Reidenbach, Zachary A. Kemmerer, Deniz Aydin, Adam Jochem, Molly T. McDevitt, Paul D. Hutchins, Emily M. Wilkerson, Jaime L. Stark, Jonathan A. Stefely, Isabel E. Johnson, Craig A. Bingman, John L. Markley, Joshua J. Coon, Matteo Dal Peraro, David J. Pagliarini

## Abstract

Human COQ8A (ADCK3) and *Saccharomyces cerevisiae* Coq8p (collectively COQ8) are UbiB family proteins essential for mitochondrial coenzyme Q (CoQ) biosynthesis. However, the biochemical activity of COQ8 and its direct role in CoQ production remain unclear, in part due to lack of known endogenous regulators of COQ8 function and of effective small molecules for probing its activity *in vivo*. Here we demonstrate that COQ8 possesses evolutionarily conserved ATPase activity that is activated by binding to membranes containing cardiolipin and by phenolic compounds that resemble CoQ pathway intermediates. We further create an analog-sensitive version of Coq8p and reveal that acute chemical inhibition of its endogenous activity in yeast is sufficient to cause respiratory deficiency concomitant with CoQ depletion. Collectively, this work defines lipid and small molecule modulators of an ancient family of atypical kinase-like proteins and establishes a chemical genetic system for further exploring the mechanistic role of COQ8 in CoQ biosynthesis.

## Introduction

The UbiB family of atypical protein kinase-like (PKL) genes is conserved across all three domains of life (Leonard et al., 1998). In eukaryotes, UbiB proteins are found exclusively in mitochondria and plastids where they have been linked to diverse processes involving cell migration, tumor cell viability, and lipid metabolism (Lundquist et al., 2012; Simpson et al., 2008; Wiedemeyer et al., 2010). An orthologous subset of UbiB proteins, including *Escherichia coli* UbiB, *S. cerevisiae* Coq8p, and human COQ8A (ADCK3) and COQ8B (ADCK4), support the biosynthesis of coenzyme Q (ubiquinone, CoQ) (Do et al., 2001; Poon et al., 2000), and mutations in COQ8A and COQ8B give rise to CoQ-related neurologic and kidney disorders, respectively (Ashraf et al., 2013; Horvath et al., 2012; Lagier-Tourenne et al., 2008; Mollet et al., 2008; Stefely et al., 2016). Despite these connections, the specific biochemical activities, regulation, and direct roles of UbiB proteins in cellular processes remain largely unclear, in part because effective tools for addressing these questions have been lacking.

To date, the only biochemical activities observed for UbiB family proteins are COQ8 *cis* autophosphorylation, which is seemingly spurious, and a low-level COQ8 ATPase activity (Stefely et al., 2016; Stefely et al., 2015). Additionally, structural analyses revealed that the COQ8 active site is likely sterically inaccessible to proteinaceous substrates, suggesting that this family possess unorthodox kinase functionality in lieu of canonical protein kinase activity. However, many members of the protein kinase-like (PKL) superfamily are subject to activation by lipids and small molecules, including AMPK (AMP) (Ferrer et al., 1985), PKA (cAMP) (Walsh et al., 1968), PKC (diacylglycerol) (Takai et al., 1979a; Takai et al., 1979b), and AKT (phosphoinositides) (Burgering and Coffer, 1995; Franke et al., 1995), indicating that COQ8 activity may be altered or enhanced in a similar manner. Consistently, we recently found that Coq8p copurifies with intermediates in the CoQ pathway (Stefely et al., 2016), and have shown that the single-pass transmembrane domain of COQ8A can induce dimerization (Khadria et al., 2014), together suggesting that COQ8 activity may particularly be regulated by interactions at the membrane; however, this remains unexplored.

A second limitation for defining the functions of UbiB proteins has been the lack of specific inhibitors that can be leveraged to disrupt protein activity acutely and reversibly. While the use of active site point mutants of COQ8 in growth assays have implied the importance of COQ8 catalytic activity for CoQ production (Stefely et al., 2015; Xie et al., 2011), this has not been tested directly by chemical inhibition *in vivo.* Chemical-genetic strategies have proven effective for kinase inhibition when specific inhibitors for the wild type kinase are lacking. This approach involves mutating a “gatekeeper” residue in the ATP binding pocket to create a larger space capable of accommodating custom, cell-permeable inhibitors that are too bulky to bind to other kinases (Bishop et al., 2000). When successful, this technique enables acute and highly specific inhibition of kinase activity *in vivo,* thereby overcoming the caveats associated with long-term adaptions that arise from genetic manipulation. While this approach has been used for various classes of kinases (Coudreuse and Nurse, 2010; Ferreira-Cerca et al., 2012; Oh et al., 2007; Rodriguez-Molina et al., 2016), it has never been attempted for the much more divergent and atypical UbiB family, nor for a mitochondria-localized kinase in general.

To address these limitations in investigating the UbiB family, we combined biophysical, biochemical, computational, and chemical-genetic analyses to explore the activity and regulation of COQ8. We found that ATPase activity is an evolutionarily conserved feature of COQ orthologs from *E. coli* to humans that is enhanced by phenolic compounds resembling CoQ biosynthetic precursors. Remarkably, we also find that mature COQ8 specifically associates with, and is activated by, cardiolipin-containing liposomes, suggesting a potential regulatory connection between mitochondrial cardiolipin and CoQ metabolism. Finally, we establish an analog-sensitive version of COQ8 in yeast and use this protein to show that inhibition of endogenous COQ8 activity is sufficient to disrupt CoQ production and cause respiratory deficiency. Overall, our work reveals lipid and small molecule modulators of this ancient kinase-like protein, and establishes a chemical-genetic tool for the further exploration of its direct role in CoQ biosynthesis.

## Results

### UbiB family ATPase activity is enhanced by phenols

To begin exploring the ability of UbiB proteins to bind lipids and small molecules, we purified a maltose binding protein (MBP)-tagged version of *E. coli* UbiB that lacks its predicted C-terminal transmembrane (TM) domains (UbiB^CΔ47^) and analyzed copurifying lipids by mass spectrometry (Figure S1A). We found that MBP-UbiB^CΔ47^ preferentially copurifies with *E. coli* CoQ biosynthesis intermediates octaprenylhydroxybenzoate (OHB) and octaprenylphenol (OPP) and that this copurification depends on the integrity of active site residues (Figures 1A, S1B–E, and Table S1). These data are strikingly similar to our previous lipid-binding data for Coq8p, indicating that the binding of CoQ precursors to UbiB family members is conserved from bacteria to eukaryotes (Stefely et al., 2016).

**Figure 1.**
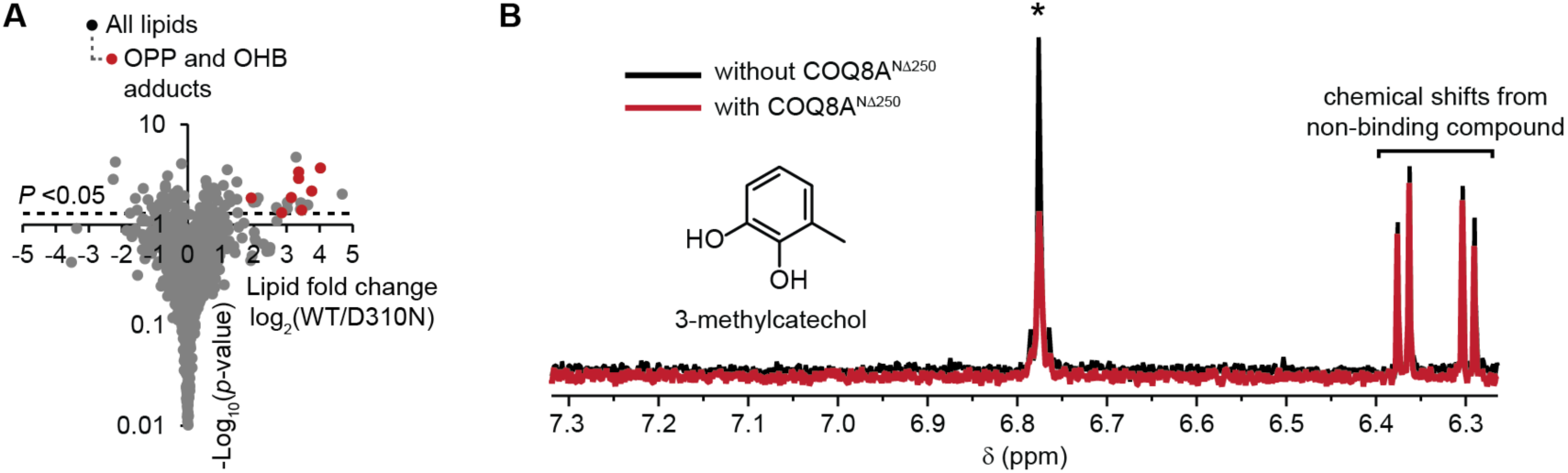
UbiB family members bind CoQ precursor-like lipids and small molecules. (A) Fold changes in the abundances of lipids copurifying with UbiB protein variants (log_2_[MBP-UbiB^CΔ47^/ MBP-UbiB^CΔ47^ D310N], n = 3) as quantified by LC-MS/MS versus statistical significance. (B) A selected region of the ^1^H-NMR spectra for a mixture containing 3-methylcatechol (3-MC) with and without COQ8A^NΔ250^. Asterisk (*) indicates a 3-MC peak that exhibits line-broadening. The peaks shown around 6.3 - 6.4 ppm belong to another compound (*cis*-3-chloroacrylic acid) in the mixture that is not interacting with COQ8A^NΔ250^.

As an orthogonal approach to assessing protein-small molecule interactions, we next performed a one-dimensional (1D) ^1^H NMR ligand-affinity screen of a diverse 418 compound library using COQ8A^NΔ250^ (COQ8A lacking its mitochondrial localization sequence and TM domain). To do so, we introduced COQ8A^NΔ250^ to mixtures of 3-5 compounds and analyzed the resulting line broadening and/or chemical shift changes of the ^1^H peaks for compounds interacting with COQ8A^NΔ250^. In addition to its expected binding to adenosine, COQ8A^NΔ250^ interacted with a total of 25 compounds including multiple that, like OPP and OHB, are phenol derivatives (Figures 1B and S1F). (Figures 1B and S1F and Table S3). Collectively, these data establish that COQ8 possesses an evolutionarily conserved capacity to bind CoQ precursor-like molecules, suggesting that these molecules are potential regulators or direct substrates of COQ8.

The CoQ precursors that copurify with Coq8p and UbiB are difficult to synthesize and are not commercially available; as such, to explore the functional ramifications of these COQ8-ligand interactions, we opted to screen a variety of CoQ headgroup-like analogs (phenols and hydroquinones) for their effect on COQ8 activity. To monitor ATP consuming activity, we used the ADP-Glo assay, which measures ADP produced as a proxy for phosphoryl transfer activity. We included the nonionic detergent Triton X-100 in our screens, as it has been shown to activate certain membrane bound or membrane associated proteins (Kanoh et al., 1991; Yoon and Robyt, 2005). Strikingly, we found that 2-alkylphenols (2-allylphenol and 2-propylphenol), which are structurally similar to OPP, strongly activated UbiB family members when in the presence of Triton X-100 (Figures 2A and S2A). This increase in activity was seen both for wild type (WT) and A339G (A-to-G) nucleotide-binding loop (P-loop) mutant, which has a higher baseline ATPase and *cis* autophosphorylation activity likely due to enhanced ATP binding (Stefely et al., 2016) (See Table S1 for explanations of mutations used in this study). The negative control D507N (D-to-N) mutant, which lacks ATP binding ability, showed no activity (Figures 2A and S2A).

**Figure 2.**
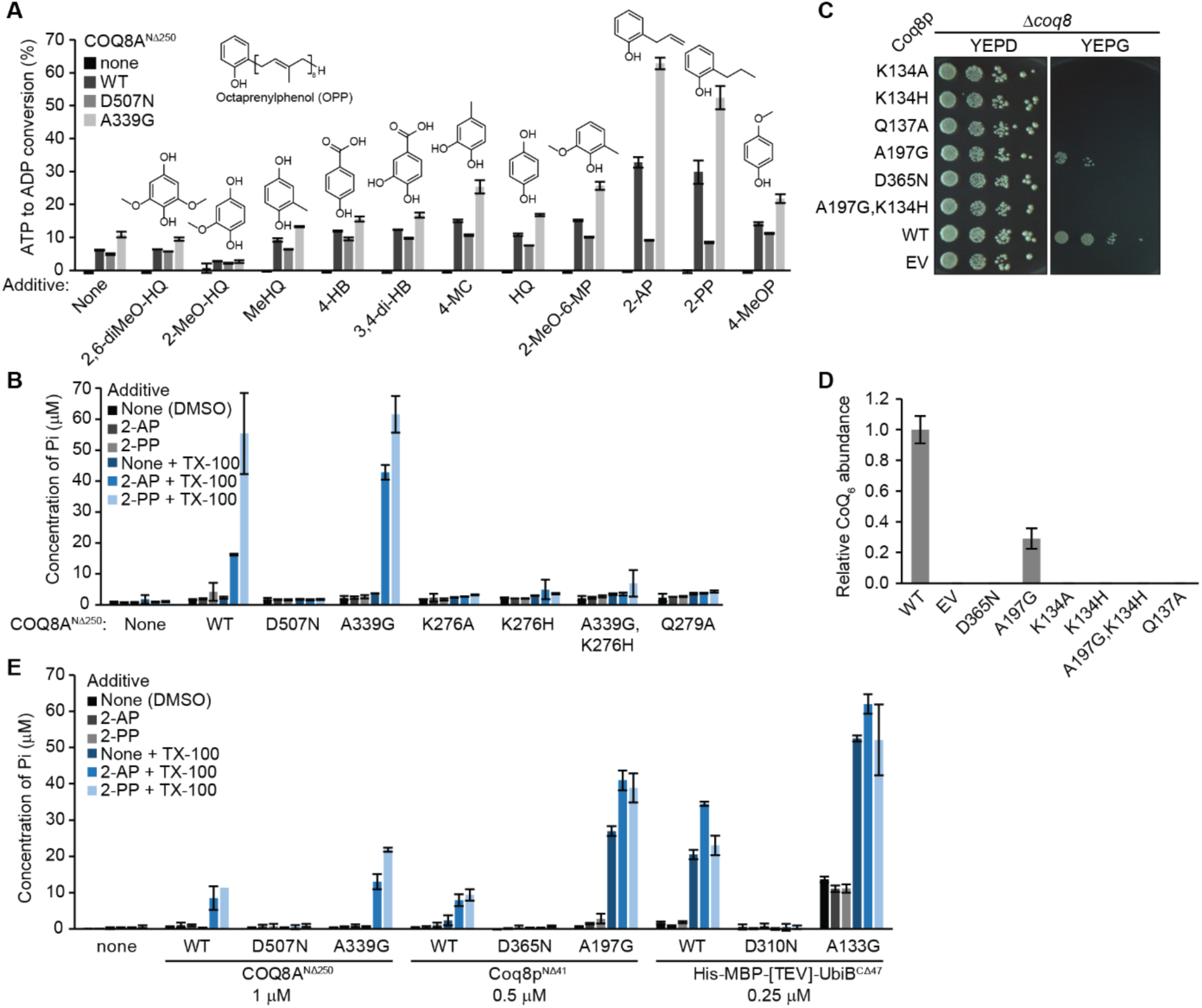
UbiB family members are activated by Triton X-100 and 2-alkylphenols. (A) Screen of CoQ intermediate-like compounds with Triton X-100 in an ADP-Glo assay. 2,6-diMeO-HQ, 2,6-dimethoxyhydroquinone; 2-MeO-HQ, 2-methoxyhydroquinone; MeHQ, methylhydroquinone; 4-HB, 4-hydroxybenzoic acid; 3,4-di-HB, 3,4-dihydroxybenzoic acid; 4-MC, 4-methylcatechol; HQ, hydroquinone; 2-MeO-6-MP, 2-methoxy-6-methylphenol; 2-AP, 2-allylphenol; 2-PP, 2-propylphenol; 4-MeOP, 4-methoxyphenol. (B) Malachite green ATPase assay with COQ8A^NΔ250^ KxGQ mutants, Triton X-100, and 2-alkylphenols. Pi, inorganic phosphate.(C) Drop assay of *Δcoq8* transformed with KxGQ mutants. (D) Relative CoQ abundance from *Δcoq8* transformed with Coq8p variants. (E) Malachite green ATPase assay of UbiB family members from human, yeast, and *E. coli* with 2-alkylphenols and Triton X-100. Concentration of protein used is listed under the protein name, and reactions were incubated at 30 °C for 10 minutes rather than 45 minutes.

Our previous structure of COQ8 revealed the presence of a water molecule in the active site bound by the glutamine (Q) of the invariant KxGQ motif that could serve as the nucleophile for an ATPase reaction (Stefely et al., 2015). As such, we hypothesized that the majority of our observed ADP production was due to ATP hydrolysis with water, which could be detected by the production of free phosphate using a malachite green assay. Indeed, we found a clear correlation between the amount of activity observed in the ADP-Glo assay and the malachite green assay, demonstrating that 2-alkylphenols with Triton X-100 increase the ATPase activity of UbiB family members (Figures 2B and S2B). Consistently, COQ8 KxGQ domain mutations that possess markedly enhanced *cis* autophosphorylation (e.g., the double A-to-G, K-to-H mutant) (Stefely et al., 2016; Stefely et al., 2015) show little ATPase activity, are unable grow in respiratory media, and fail to enable CoQ_6_ production *in vivo* (Figures 2C, 2D, S2C, and S2D). This further demonstrates that ATPase activity, and not protein kinase-related activity, tracks most closely with the endogenous function of COQ8.

Notably, compounds that more closely mimic the mature CoQ headgroup, such as 2,6-dimethoxyhydroquinone or CoQ_1_, did not greatly enhance COQ8 activity, in agreement with our CoQ precursor binding data (Figure S2E) (Stefely et al., 2016). Furthermore, the activation by 2-alkylphenols with Triton X-100 is conserved in UbiB, Coq8p, and COQ8A, mirroring the conservation of CoQ precursor binding (Figure 2E). Overall, these data suggest that COQ8 possesses an evolutionarily conserved ability to bind early CoQ intermediates and to be activated by 2-alkylphenols.

### Mature COQ8 is activated by liposomes

Endogenous COQ8 contains a TM domain that anchors it to the inner mitochondrial membrane facing the matrix where the other complex Q proteins reside and where it can potentially interact with CoQ intermediates (He et al., 2014; Tauche et al., 2008). This suggests that membrane association might be another important modulator of its activity, as has been shown for other membrane-bound kinases (Jura et al., 2009). The mature N-termini of COQ8A and Coq8p (i.e., the N-termini present after cleavage of the mitochondrial localization sequences) begin 163 residues and 42 residues from the unprocessed N-terminus, respectively (Calvo et al., 2017; Cullen et al., 2016; Stefely et al., 2015; Vogtle et al., 2009). To test the effect of an intact TM domain on COQ8 membrane association and activity, we purified recombinant versions of the mature forms of COQ8A (COQ8A^NΔ162^) and Coq8p (Coq8p^NΔ41^), marking the first time that a mature form of a human UbiB family member has been purified (Figure S3A).

Given that mature COQ8 resides on the inner mitochondrial membrane (IMM), we reasoned that mixing the protein with liposomes that contain IMM-enriched lipids, such as cardiolipin (CL) and phosphatidylethanolamine (PE) (de Kroon et al., 1997; Horvath and Daum, 2013; Zinser et al., 1991), along with the abundant membrane lipid phosphatidylcholine (PC), could have an effect on COQ8 membrane binding and activity. To test this, we first used a liposome flotation assay to determine how effectively COQ8 binds to liposomes (Figure 3A). Without the TM domain, COQ8A^NΔ250^ exhibited moderate intrinsic affinity for PC/PE/CL liposomes; however, the addition of the TM domain enabled near complete COQ8A^NΔ162^ and Coq8p^NΔ41^ liposome binding (Figures 3B). Strikingly, this association markedly enhanced the ATPase activity of COQ8A^NΔ162^ and Coq8p^NΔ41^ (Figure 3C). These data imply that UbiB proteins, like other PKL family members, may be activated endogenously by binding to particular membrane environments.

**Figure 3.**
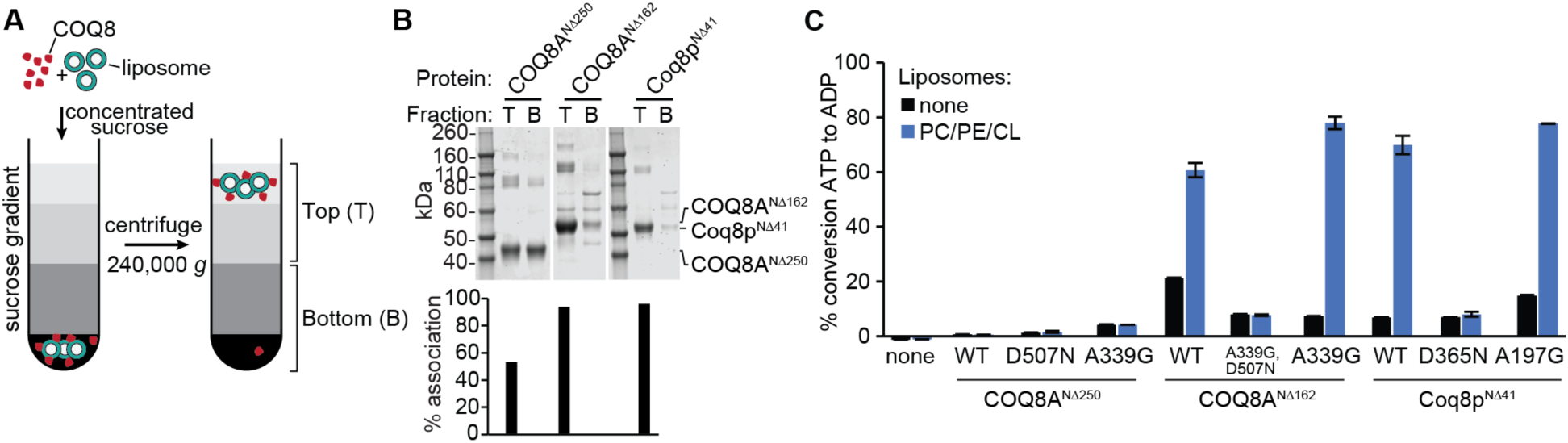
COQ8 binds to liposomes. (A) Schematic of a liposome flotation assay. (B) Coomassie stained SDS-PAGE from a liposome flotation assay with COQ8 and PC/PE/CL liposomes. T; top fraction, B, bottom fraction. (C) ADPGlo assay with COQ8 variants and PC/PE/CL liposomes.

### Cardiolipin specifically increases COQ8 membrane interaction and ATPase activity

To determine whether the COQ8 membrane binding and activity is driven by a particular membrane component, we tested the ability of multiple individual lipids to activate COQ8 when reconstituted in PC carrier liposomes. Remarkably, from amongst the eight individual species tested, only CL was able to activate COQ8 (Figure 4A). This increase in activity was matched by the superior ability of CL to facilitate binding of COQ8 to liposomes, indicating that CL activation of COQ8 is directly linked to its ability to mediate COQ8-membrane interactions (Figures 4B and S4A). Interestingly, COQ8 lacking its TM still preferentially bound to CL-containing liposomes (Figure S4B), but was not activated by them (Figure 3C), suggesting that the TM domain itself is a regulatory feature of COQ8.

To test whether liposome/CL binding specifically enhances the ATPase activity of COQ8, we performed [γ-^32^P]ATP kinase reactions with liposomes to follow the location of the gamma phosphate. We also assessed autophosphorylation using SDS-PAGE and potential phosphorylation of copurifying lipid substrates or CL using thin-layer chromatography. Consistent with the activation of COQ8 by phenols, no autophosphorylation or lipid phosphorylation was observed (data not shown), further supporting the hypothesis that COQ8 acts as an ATPase (Figures S4C). Additionally, liposome-activation of COQ8 did not alter its inability to phosphorylate purified protein substrates, including complex Q proteins and myelin basic protein (Figures S4D and S4E). Collectively, these data demonstrate that CL enables COQ8 membrane binding and enhances COQ8 ATPase activity.

**Figure 4.**
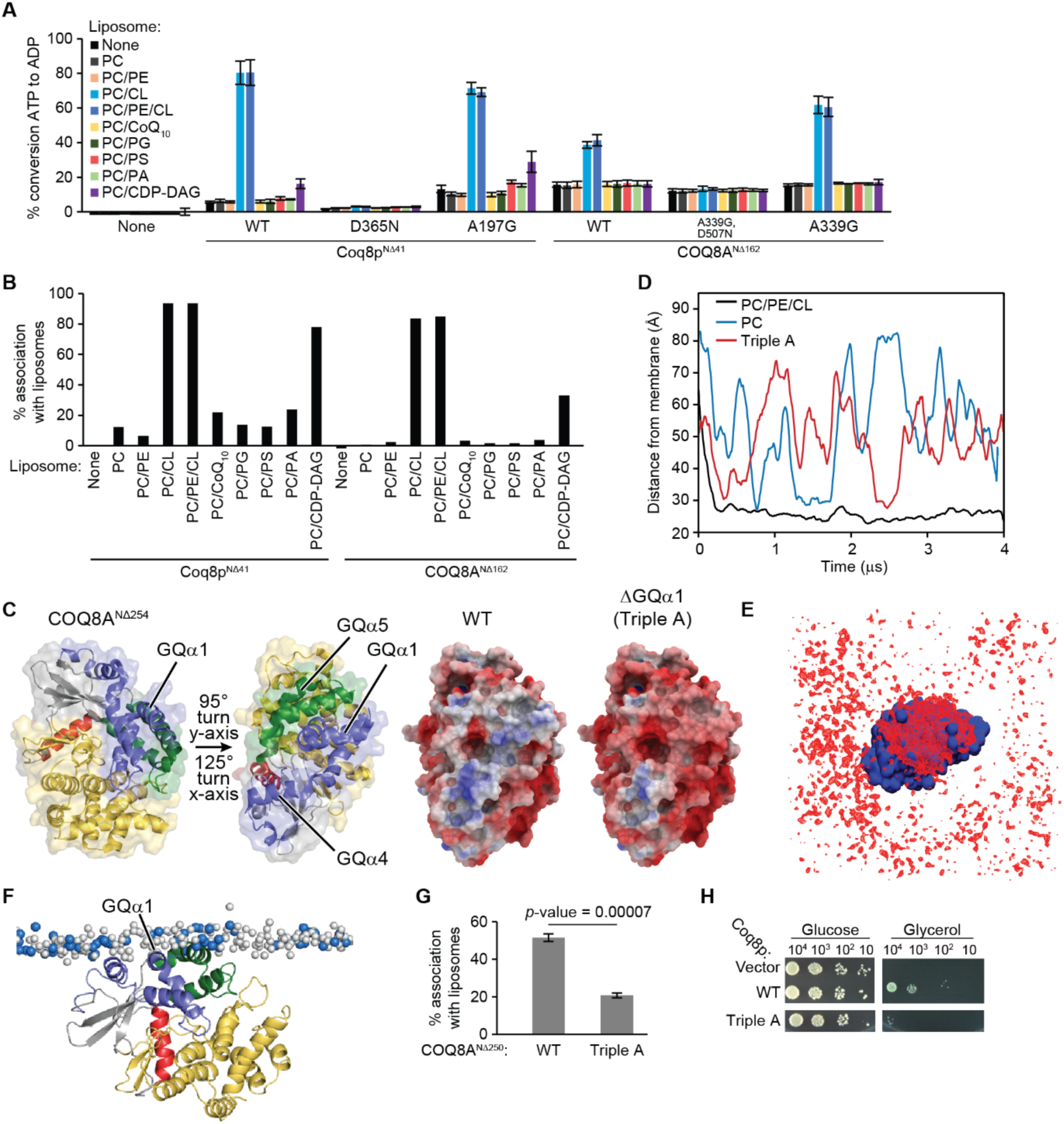
CL enhances the ATPase activity of COQ8 and liposome binding. (A) ADP-Glo assay with a panel of liposomes and TM domain containing COQ8. Error bars represent s.d. of two independent experiments performed in technical duplicate. (B) Liposome flotation assay with a panel of PC based liposomes and TM-containing COQ8. (C) The two structures of COQ8A on the left are colored according to Stefely et al. (2015) and turned to show the orientation of the electrostatic maps (Stefely et al., 2015). The KxGQ domain is colored in purple and green in the structures on the left. On the right, there are electrostatic maps of COQ8A showing the positively charged region spanning GQα1, GQα4, and GQα5 [negative (−5 kcal/e.u. charge): red, via white, to positive (+5 kcal/e.u. charge): blue]. (D) Time evolution of the distance between the center of mass of the protein and the center of mass of the phosphate heads of the leaflet it interacts with for CG-MD simulations of COQ8A with PC or PC/PE/CL liposomes or of the Triple A mutant with PC\PE\CL liposomes. (E) Average occupancy of CL phosphate heads (red) when COQ8A (blue) is centered throughout the trajectory. (F) Snapshot from a CG-MD simulation (t ~ 4 μs) showing the interaction of COQ8A with a PC/PE/CL membrane. CL phosphate heads shown in blue and PC/PE phosphate heads shown in white. (G) Percent protein associating with liposomes from a liposome flotation assay with COQ8A^NΔ250^ WT and the Triple A mutant (R262A,R265A,K269A). (H) Serial dilutions of Δ*coq8* transformed with Coq8p WT or the Triple A (R120A,K124A,K127A) on synthetic complete glucose (2%) or glycerol (3%) containing media.

### COQ8 interacts with the membrane through its signature KxGQ domain

To further explore the nature of COQ8 membrane binding and activation, we used molecular dynamics to simulate the binding of COQ8A to membranes. Examination of the electrostatic surface potential of the soluble domain of COQ8A revealed an extensive positive patch generated primarily by the GQα1 and GQα4 helices that likely interacts with negatively charged phospholipids found in the IMM, such as the CL used in our liposomes (Figures 4C and S4F). We then used coarse-grained molecular dynamics (CG-MD) simulations to investigate the binding of COQ8A to PC/PE/CL, mimicking the IMM-like membranes used in our liposome flotation assays, or to PC alone, mimicking a non-IMM membrane model. Invariably, the presence of CL at physiological concentration (Horvath and Daum, 2013) was required to induce rapid (within the first microsecond) membrane association (Figure 4D and Movies S1 and S2) mediated by the GQα1 and GQα4 helices (Figures 4F), providing independent validation of our liposome flotation assay results. Interestingly, anchoring of COQ8A at the IMM is associated with local reorganization of lipids, featuring a strong segregation of CL species to the COQ8A interface (Figure 4E).

Our CG-MD simulations further predicted three positively charged residues of the GQα1 helix (R262, R265, and K269) as the key anchor points driving initial encounter of the soluble domain with the membrane via electrostatic interactions with cardiolipin phosphate heads (Figures S4G and S4H). To validate the importance of these residues, we purified a triple mutant of COQ8A^NΔ250^ (R262A,R265A,K269A; Triple A; ΔGQα1) and measured its ability to bind liposomes. Indeed, this mutant demonstrated a large decrease in liposome association (Figures 4G and S4I). When the corresponding mutations were introduced into *coq8,* its ability to rescue Δ*coq8* respiratory deficiency was significantly diminished, suggesting that these residues may have important ramifications for membrane binding and/or orientation *in vivo* (Figures 4H and S4J). Altogether, these simulations reinforce the importance of CL for COQ8 membrane association, and suggest that the COQ8–CL interaction is driven by conserved residues in the signature KxGQ domain.

### Inhibition of Coq8p-*AS1* activity disrupts CoQ biosynthesis

Our analyses above suggest that COQ8 is a conserved ATPase that is activated by lipids found in its native environment on the mitochondrial inner membrane. We next sought to determine whether acute chemical inhibition of COQ8 activity in this native environment would be sufficient to disrupt CoQ biosynthesis and respiratory function in yeast. To do so, we attempted to generate an analog-sensitive (*AS*) version of Coq8p (Coq8p-*AS*1) using a chemical-genetic approach (Bishop et al., 2000).

To create Coq8p-*AS*1, we first attempted to identify its putative gatekeeper residue by performing sequence and structural alignments of Coq8p with Src, a kinase known to be amenable to chemical-genetic modulation (Figures 5A and 5B) (Zhang et al., 2013). These analyses suggested that methionine 303 (M303) is the Coq8p gatekeeper residue, which we further tested *in silico.* To do so, we changed M303 to a glycine residue and modeled the binding of the custom chemical-genetic inhibitor, 3-MB-PP1 (Bishop et al., 1999; Burkard et al., 2007). The WT Coq8p nucleotide binding pocket was incapable of accepting 3-MB-PP1 due to steric clashes with the methionine side chain (Figure 5C); however, the M303G model predicted that it would be able to accommodate 3-MB-PP1 (Figures 5D and 5E).

**Figure 5.**
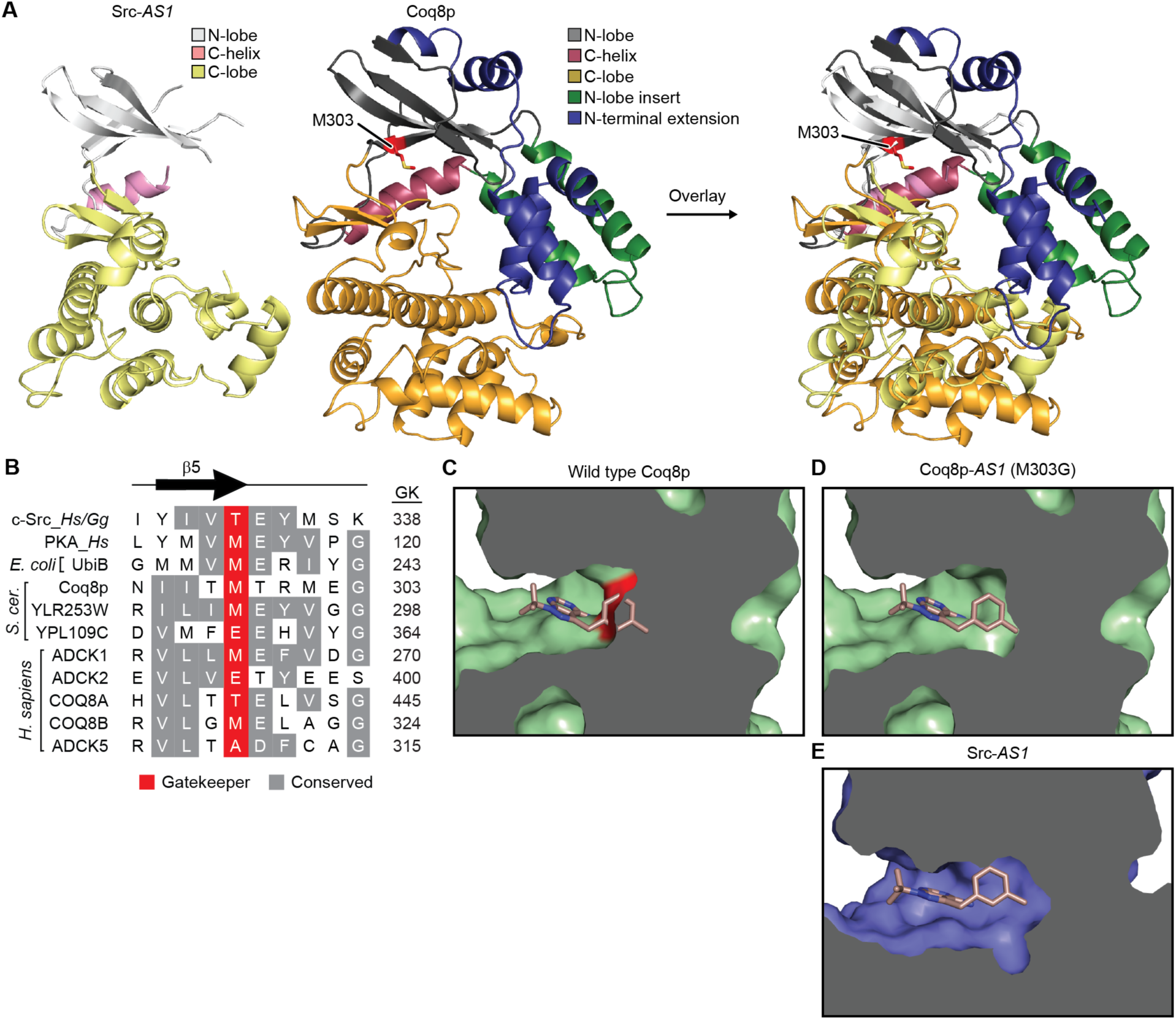
Creation of analog-sensitive Coq8p. (A) Structural alignment of Coq8p homology model based on COQ8A (PDB: 5I35) with Src-*AS*1 (PDB: 4LGG). The M303 gatekeeper of Coq8p is shown in red. (B) Primary sequence alignment of the gatekeeper residues from Src, PKA, and the UbiB family. Hs, *Homo sapiens*; Gg, *Gallus gallus*; GK, gate keeper residue number. (C) Modeling of 3-MB-PP1 in the active site of the homology model of Coq8p. The surface of M303 is colored in red. (D) Mutation of methionine 303 to a glycine (M303G) creates a binding pocket for 3-MB-PP1. (E) Structure of Src-*AS*1 with 3-MB-PP1 bound from PDB: 4LGG (Zhang et al., 2013).

To generate this putative Coq8p-*AS*1, we expressed and purified WT and M303G Coq8p^NΔ41^ from *E. coli* (Figure S5A). Using differential scanning fluorimetry (Niesen et al., 2007), we tested the predicted ability of Coq8p^NΔ41^ M303G to bind nucleotides and inhibitors designed specifically for *AS-*kinases. Consistent with the modeling predictions, only the M303G mutant was able to bind to the inhibitor 3-MB-PP1, which increased the melting temperature of the mutant by a considerable nine degrees (Figures 6A and 6B). Similar to other *AS-*kinases, this gatekeeper mutant bound nucleotides with a decreased apparent dissociation constant (*K*_d, app_) compared to the WT enzyme (Figure S6A) (Bishop et al., 2000; Zhang et al., 2005). Nonetheless, the M303G mutant was able to rescue Δ*coq8* respiratory function (Figure 6C).

**Figure 6.**
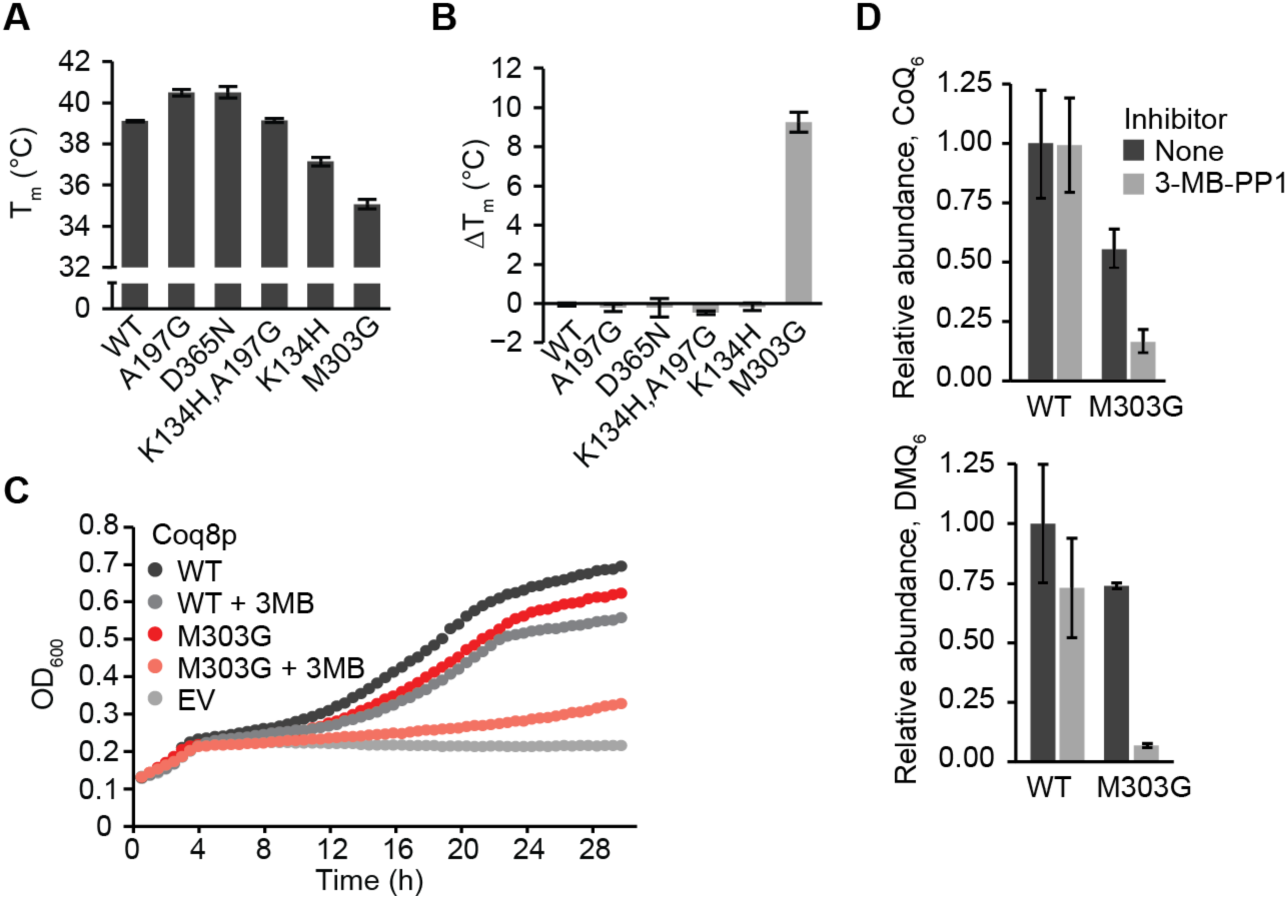
Acute inhibition of Coq8p decreases CoQ biosynthesis. (A) Melting temperature of Coq8p variants measured with DSF. (B) ΔT_m_ for COQ8 variants with the *AS-*inhibitor 3-MB-PP1. (C) Growth curve of Δ*coq8* yeast transformed with Coq8p-*AS*1 and treated with 3-MB-PP1 (25 μM) in Ura^−^, pABA^−^ 0.1% glucose 3% glycerol media. 3MB, 3-MB-PP1; EV, empty vector. (D) Relative abundance of CoQ_6_ and DMQ_6_ from Δ*coq8* yeast transformed with Coq8p-*AS*1 and treated with 3-MB-PP1 (25 μM) after 24 hours measured using LC-MS/MS.

To determine the effects of endogenous Coq8p inhibition, we treated Δ*coq8* yeast expressing Coq8p M303G with 3-MB-PP1 and measured respiratory function and the production of CoQ and the CoQ precursor demethoxy-coenzyme Q (DMQ) at 24 hours after inhibitor addition (Figure S6B). Treatment of Coq8p-*AS*1 yeast with 3-MB-PP1 caused a marked decrease in respiratory function as assessed by severely delayed growth assay in liquid culture (Figure 6C). Furthermore, this respiratory deficiency was accompanied by disrupted CoQ biosynthesis. In particular, levels of CoQ and, even more so, DMQ were diminished 24 hours after inhibitor treatment (Figure 6D). These data reveal that 3-MB-PP1 added to yeast liquid culture is capable of accessing and inhibiting Coq8p-*AS*1 in its proper endogenous location within the mitochondrial matrix, and that COQ8 catalytic activity is required for CoQ precursor processing upstream of DMQ, consistent with earlier analyses of Δ*coq8* yeast and of strains harboring *coq8* mutations. Collectively, our work defines lipid and small molecule modulators — both endogenous and custom *AS-*compounds — of an ancient atypical kinase-like protein. Additionally, our establishment of a chemical genetic system for acutely inactivating COQ8 will enable further dissection of the precise mechanistic role of COQ8 in CoQ biosynthesis.

## Discussion

Coenzyme Q was discovered 60 years ago as a key component of the mitochondrial electron transport chain, yet multiple aspects of its biosynthesis remain obscure. One such aspect is the unexplained need for multiple auxiliary proteins that have no clear catalytic role in the pathway, including COQ8 — a highly conserved member of the large protein kinase-like (PKL) superfamily. Given its PKL membership, COQ8 has been proposed to fill a regulatory gap in CoQ biosynthesis by phosphorylating other proteins in the CoQ pathway (Xie et al., 2011). However, to date, no COQ8 *trans* protein kinase activity has been observed, demanding that other models for its activity be explored.

To further investigate the biochemical function of COQ8, we sought to study it in its native environment and to establish new tools to modulate its function. Multiple members of the PKL superfamily become activated by binding to endogenous molecules found at their sites of cellular activity, such as the activation of PKC or Akt by diacylglycerol and phosphoinositide 3,4,5-triphosphate, respectively, at the plasma membrane (Burgering and Coffer, 1995; Franke et al., 1995; Nishizuka, 1992). Furthermore, protein-lipid interactions in general are becoming increasingly recognized as regulatory and structural features of proteins (Gupta et al., 2017; Moravcevic et al., 2012; Yeagle, 2014), indicating that it may be important to identify the full set of molecules that interact with COQ8 in order to elucidate its biochemical function.

Consistent with our recent results using Coq8p, our binding assays point to a conserved and selective interaction between COQ8 and precursors in the CoQ biosynthesis pathway. Curiously, these interactions seem to rely on the integrity of the COQ8 active site and result in an increase in COQ8 ATPase activity without affecting its inability to phosphorylate proteinaceous substrates. In mitochondria, these precursors would most likely be found buried between the leaflets of the inner mitochondrial membrane (IMM) due to their extreme hydrophobicity. These data might suggest that COQ8 is poised to sense CoQ precursors within the membrane and to then couple the hydrolysis of ATP to the partial extraction of these lipids into the aqueous matrix environment where they could be modified by other CoQ biosynthesis enzymes. Testing this model directly will require the chemical synthesis of CoQ precursor molecules (Barkovich et al., 1997; van der Klei et al., 2002) or their purification from endogenous sources like Δ*ubi E. coli* (Aussel et al., 2014) or Δ*coq S. cerevisiae* overexpressing Coq8p (Xie et al., 2012). Alternatively, it remains possible that the observed ATPase activity is a proxy for small molecule kinase activity *in vivo*; however, as COQ8-dependent small molecule phosphorylation was not observed in our assays, these data indicate that endogenous small molecule substrate(s) would likely be distinct from the free hydroxyl group(s) of CoQ intermediates, the most intuitive candidate substrates.

Similar to PKC and Akt, our data also indicate that COQ8 is activated via interaction with a specific lipid — cardiolipin (CL). CL is a unique anionic, four-tailed lipid found at particularly high concentrations in the IMM where it influences the activity and stability of various mitochondrial membrane proteins (Claypool, 2009; Claypool et al., 2008; Gebert et al., 2009; Planas-Iglesias et al., 2015), and promotes the formation of respiratory chain supercomplexes (Pfeiffer et al., 2003). The specific interaction of COQ8 with CL is consistent with COQ8’s localization to the IMM (Cullen et al., 2016; He et al., 2014; Tauche et al., 2008), where the other complex Q proteins reside. In addition to mere binding, our CG-MD simulations indicate that CL also directs the orientation of the soluble COQ8 domain along the membrane surface, perhaps thereby enabling COQ8’s known interactions with other COQ proteins (Cullen et al., 2016; Floyd et al., 2016) and/or with membrane-embedded CoQ precursors. Furthermore, CL is thought to be distributed non-uniformly throughout the IMM (Mileykovskaya and Dowhan, 2009), suggesting that the interaction of COQ8 with CL might seed complex Q formation at strategic sub-mitochondrial. These observations suggest models whereby the proper positioning of COQ8 at CL-rich domains on the mitochondrial inner membrane activates its ATPase activity as a requisite first step in enabling complex Q assembly and/or CoQ biosynthesis, and that exposure to CoQ intermediates at those sites can further enhance COQ8 function (Figure 7). Within this context, CG-MD shows once more to be a powerful resource to understand the molecular determinants underlying protein-lipid interplay.

**Figure 7.**
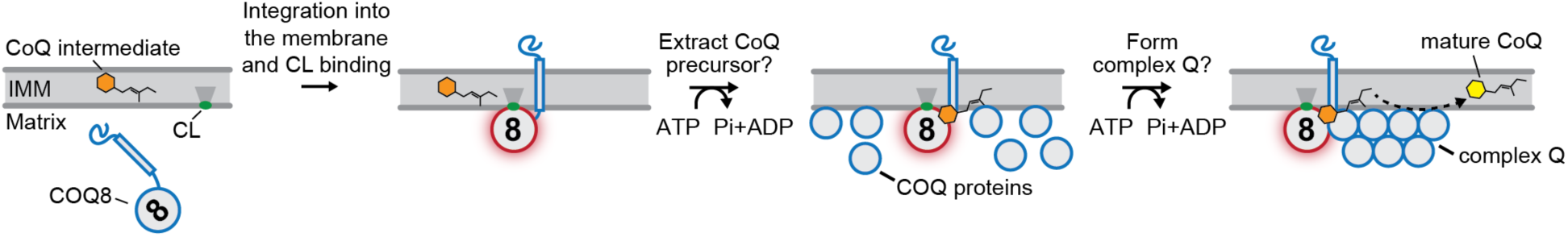
Model for COQ8 ATPase ability could facilitate CoQ biosynthesis. First, COQ8 is imported in to the matrix. Binding to CL then facilitates COQ8’s association with, and insertion into, the IMM with its soluble domain properly oriented along the membrane surface. Next, the CL-induced activation of COQ8 (red glow) allows it to advance CoQ biosynthesis by coupling the hydrolysis of ATP to the extraction of CoQ precursors out of the IMM and/or to the formation of complex Q.

More broadly, these results suggest an important and underappreciated connection between CoQ and CL — two quintessential mitochondrial lipids that enable oxidative phosphorylation (OxPhos). Indeed, CL has long been known to enable electron transfer between CI and CIII (Fry and Green, 1981), and in plants, CL was found to have a positive effect on the electron transfer rate from CoQ to the photosynthetic reaction center (Giustini et al., 2005). Similarly, CL enhances electron transport from Complex I to CoQ in *Drosophila* mitochondrial lysate (Vos et al., 2017), and the CL remodeling phospholipase Cld1p was shown to rescue *coq7* hypomorphs in *S. cerevisiae* harboring mutations in Coq7p’s hydrophobic α-helix that is predicted to bind the IMM (Kar et al., 2016). These observations in conjunction with our data suggest a model whereby enhanced CL biosynthesis may augment OxPhos function in part by activating the biosynthesis of CoQ via COQ8.

Testing these and other models of COQ8 function require new tools to modulate COQ8 activity in its endogenous setting where these activating molecules are present. To that end, we developed an analog-sensitive (*AS*) version of yeast Coq8p (Coq8p-*AS*1). This chemical-genetic strategy has proven to be highly effective for elucidating the functions of other PKL family members (Coudreuse and Nurse, 2010; Ferreira-Cerca et al., 2012; Oh et al., 2007; Rodriguez-Molina et al., 2016), but has never been attempted for the highly divergent UbiB family. Furthermore, this approach has not been attempted for kinases that reside within organelles where they would potentially be inaccessible to inhibitors. Our work demonstrates that endogenous chemical inhibition of COQ8 is sufficient to disrupt CoQ biosynthesis, further suggesting that the catalytic activity of COQ8 is important for this pathway. Moreover, our demonstration that a UbiB family member is amenable to a chemical genetic system suggests that this tool can likewise be used to interrogate the functions of other family members, including Ypl109c and Ylr253w (Mcp2p) in yeast, the ABC1Ks in plants, and ADCK1,2 & 5 in humans. These proteins have been implicated in diverse biological functions in cell migration (Simpson et al., 2008), tumor cell viability (Brough et al., 2011; Wiedemeyer et al., 2010), and lipid metabolism (Lundquist et al., 2012; Martinis et al., 2013; Tan et al., 2013), each through unclear means. A chemical genetic approach would offer the opportunity to identify functions for these proteins without the added complexity of compensatory changes that accompany full gene knockouts.

### Significance

UbiB proteins comprise a large and ancient family of protein kinase-like (PKL) molecules that spans all three domains of life. These proteins have been implicated in diverse cellular processes and are mutated in human diseases; however, little is known about their direct biochemical roles and activities. The main orthologous UbiB proteins investigated in this work — *S. cerevisiae* Coq8p and human COQ8A (jointly referred to as COQ8) — exemplify this problem: they are known to enable the biosynthesis of the key mitochondrial lipid, coenzyme Q (CoQ), but the mechanism is unclear. Investigations into the functions of other PKL family members have benefitted greatly from the identification of activating lipids and small molecules, and from the use of custom inhibitors, suggesting that such tools may likewise be beneficial in elucidating how UbiB proteins operate; however, such resources have not been established for this family. Our work helps to address these deficiencies by defining lipid and small molecule modulators of COQ8. We demonstrate that COQ8 exhibits a conserved ATPase activity that is activated by CoQ-like precursors and by binding to membranes that contain cardiolipin — a prevalent mitochondrial lipid enriched at the inner mitochondrial membrane (IMM) where COQ8 resides. We also establish an “analog-sensitive” version of Coq8p that can be inhibited by small molecules *in vivo* and use this tool to demonstrate that acute inhibition of Coq8p is sufficient to cause CoQ deficiency and respiratory dysfunction. While further efforts are needed to pinpoint the precise biochemical role of COQ8 in enabling CoQ biosynthesis, our work significantly advances our understanding of the core COQ8 biochemical function across evolution (ATPase activity), reveals how the positioning of COQ8 on the IMM is key to its activation, and provides effective new tools for the further investigation of the role of COQ8 in CoQ biosynthesis.

## SUPPLEMENTAL INFORMATION

Supplemental Information includes Supplemental Experimental Procedures, six Figures, two Movies, and three Tables.

## AUTHOR CONTRIBUTIONS

A.G.R. and D.J.P. conceived of the project and its design. Z.A.K., D.A., A.J., P.D.H., M.T.M., J.S., E.M.W., J.A.S., I.E.J., C.A.B., and M.D.P. performed experiments and data analysis. D.A. and M.D.P designed and performed molecular modeling and simulation that supported the experimental design. J.L.M., J.J.C., M.D.P., and D.J.P. provided key experimental resources and/or aided in experimental design. A.G.R. and D.J.P. wrote the manuscript.

## ACKNOWLEDGMENTS

We would like to thank Professor Kevan Shokat and Flora Rutaganira for useful discussions and for generously providing the 3-MB-PP1 used in this study. Research reported in this publication was supported by the National Institute of General Medical Sciences of the National Institutes of Health under award numbers R01GM112057 (to D.J.P.), T32GM008505 (to A.G.R.), T32HL007899 (to E.W), and R35GM118110 (to J.J.C). This work was further supported by SNSF grants 31003A_170154 and 200021_157217 (to M.D.P.). The content is solely the responsibility of the authors and does not necessarily represent the official views of the National Institutes of Health.

## STAR METHODS

### CONTACT FOR REAGENT AND RESOURCE SHARING

Further information and requests for resources and reagents should be directed to and will be fulfilled by the Lead Contact, Dave Pagliarini.

### EXPERIMENTAL MODEL AND SUBJECT DETAILS

*Escherichia coli* strain DH5α (NEB) was used for all cloning applications and grown at 37 °C in LB media with antibiotics. *Escherichia coli* strain BL21-CodonPlus (DE3)-RIPL (Agilent) was used for all protein expression and purification purposes. See methods below for more details. For yeast rescue assays in Figures 2C, 2D, 4H, and S4L, *Saccharomyces cerevisiae* strain W303 was used. For the analog sensitive experiments (Figure 6C and 6D), *Saccharomyces cerevisiae* strain BY4742 was used. See methods below for more details.

## METHOD DETAILS

### NMR screening

One dimensional (1D) NMR ligand-affinity screens provide a quick and effective approach to identify small molecules from a large library of compounds that interact with a protein (Fischer and Jardetzky, 1965; Lepre et al., 2004; Mercier et al., 2006). We used a 1D ^1^H NMR ligand affinity screen to identify interacting small molecules from a library of 418 compounds, which consists primarily of metabolites, substrates, inhibitors, etc (Table S3). To minimize the materials required to evaluate every compound with COQ8A^NΔ250^, the compounds were combined into 89 mixtures of 3-5 compounds each. The mixtures were designed to minimize ^1^H peak signal overlap between the compounds using NMRmix (Stark et al., 2016). Two sets of NMR samples were created for each mixture: one with COQ8A^NΔ250^ and one without COQ8A^NΔ250^. The NMR samples were prepared in 15 mM bis-Tris-d_19_ buffer at pH 7.4 with 250 μM NaCl, 0.04% NaN_3_, and 15 μM DSS in in 99.99% D_2_O and placed in 3 mm SampleJet NMR tubes. Each NMR sample had a 75 μM final concentration for each compound in the mixture and a final protein concentration of 6 μM. The NMR spectra were collected on a Bruker Avance III 600 MHz spectrometer equipped with a 5 mM cryoprobe and a SampleJet sample changer set at 25 °C, and Topspin v3 (Bruker) and NMRbot were used to automate the data collection process (Clos et al., 2013). The 1D ^1^H spectra were collected for all samples using a Carr-Purcell-Meiboom-Gill (CPMG) pulse sequence with *f*_1_ presaturation and 128 averaged transients consisting of 16,384 time-domain points and a sweep width of 12 ppm.

The 1D ^1^H spectra were phased, baseline corrected, referenced to DSS, and analyzed using MestReNova 11 (Mestrelab Research). The spectra of the mixtures with COQ8A^NΔ250^ were visually compared to the spectra of the mixtures without COQ8A^NΔ250^. If the peak intensities of a compound in a mixture showed a substantial decrease (typically >50%) in the presence of COQ8A^NΔ250^, the compound would be considered a “hit” or interacting compound. The screen of the compound library with COQ8A^NΔ250^ identified 25 interacting compounds (Table S3).

### Computational Modeling and Coarse-Grained Molecular Dynamics Simulations

All simulations were performed using the GROMACS (Pronk et al., 2013) simulation package version 5.0.4. The systems were described with the MARTINI (Marrink et al., 2007) CG force field, together with the ElNeDyn (Periole et al., 2009) approach to maintain the secondary and tertiary structure of the protein. CG-MD simulations based on the MARTINI force field have been previously used with success for different systems to investigate protein-lipid interactions (Periole et al., 2007; Schafer et al., 2011; Stansfeld et al., 2009; van den Bogaart et al., 2011).

The systems were built with center of mass of COQ8A (PDB: 4PED) 70 Å away from the surface of the membrane and checked for whether the protein approaches and interacts with the membrane. Inner mitochondrial membrane (IMM) models were built according to the experimental molecular ratio of CL observed for bovine heart mitochondria, namely 15 to 20% of the phosphorus content (Daum, 1985). Therefore, IMM mimics were featuring a lipid concentration of 16% CL/41% POPC/37% POPE/6% DSPC, while generic membrane models featuring 100% POPC lipids were built as control. Lipid bilayers of 130 × 130 Å^2^ were generated using the insane (INSert membrANE) (Wassenaar et al., 2015) method of MARTINI. Systems were solvated with polarizable CG water and counter ions were added to neutralize the system. Following solvation, systems were energy minimized with a timestep of 5 fs. Successive equilibrations with decreasing restraints were performed in order to obtain a fully equilibrated system (force constants of 1000, 500, 250 and 0 kJ/mol applied on protein particles, 10 ns of run with a timestep of 10 fs for each). In the production phase, which is 4 μs for all simulations, the protein, membrane and the aqueous phase (water and ions) were coupled independently to an external bath at 310 K and 1 bar. Three MD repeats with randomized initial velocities were performed for each membrane system and they yielded consistent results. Average occupancy of cardiolipins was calculated using VMD’s VolMap plugin (Humphrey et al., 1996).

Atomistic models of the different GQα mutated COQ8A proteins were built in PyMOL by mutating the following residues to alanine: R262, R265, K269 for ΔGQα1; K310, K314 for ΔGQα4; R262, R265, K269, K310, K314 for ΔGQα1α4; Q366, S367, N369, S370, N373, N374 for ΔGQα5. The effect of these mutations on the electrostatic potential surface of the protein was visualized through the ICM-REBEL module of the ICM program (Abagyan et al., 1994). CG-MD simulations were performed for these four mutants following the procedure explained above, where the residues listed for each mutant were altered by the removal of charges of the polarizable CG beads, to knockout Coulomb interactions without changing the residue types.

### Modeling of Coq8p-*AS*1 with 3-MB-PP1

A homology model of Coq8p-*AS*1 was generated by SWISS-MODEL (Biasini et al., 2014) using the nucleotide-bound form of COQ8A (PDB:5I35) as a template (RMSD 0.35 Å over 2264 of 2590 aligned atoms.) To understand the likely position of the bulky analog 3-MB-PP1 bound to Coq8p-*AS*1, the structure of an analog sensitive Src kinase bound to the 3-MB-PP1 (PDB:4LGG) (Zhang et al., 2013) was superposed on the homology model for Coq8p-*AS*1 (9 Å RMSD over 1061 of 1227 atoms aligned.) The adenine ring of the nucleotide bound to COQ8A was parallel to the aromatic ring structure in 3-MB-PP1. A pairwise alignment of the adenine ring and corresponding atoms in 3-MB-PP1 (RMSD of 0.067 Å over 11 atoms) was performed to give the final model of 3-MB-PP1 bound to Coq8p-*AS*1. Electrostatic surfaces were calculated using PDB2PQR (Dolinsky et al., 2007) and APBS (Baker et al., 2001) plugins in PyMOL (v1.8).

### DNA Constructs and Cloning

Cloning of COQ8A^NΔ250^, Coq8p^NΔ41^, PKA, and p426 GPD *coq8* plasmids was previously described (Stefely et al., 2016; Stefely et al., 2015). For MBP-UbiB^CΔ47^ and COQ8A^NΔ162^ standard PIPE cloning methods (Klock et al., 2008) were used. PIPE reactions were DpnI digested and transformed into DH5α competent *E. coli* cells. Plasmids were isolated from transformants and DNA sequencing was used to identify those containing the correct constructs. pVP68K, a plasmid for expression of recombinant proteins in bacteria [8His-cytoplasmically-targeted maltose-binding protein (MBP) with a linker including a tobacco etch virus (TEV) protease recognition site fused to the protein construct (8His-MBP-[TEV]-Protein)], has been described previously (Blommel et al., 2009). Oligonucleotides were purchased from IDT (Coralville, IA, USA). For yeast rescue experiments, *coq8* was cloned into the p426 GPD vector (Mumberg et al., 1995). Point mutations were introduced by PCR-based mutagenesis and confirmed by DNA sequencing.

### Recombinant Protein Expression and Purification

Human COQ8A*^NΔ162^* and COQ8A*^NΔ250^*. Method adapted from (Stefely et al., 2016; Stefely et al., 2015).

COQ8A constructs were overexpressed in *E. coli* by autoinduction (Fox and Blommel, 2009). Cells were isolated and frozen at −80 °C until further use. For protein purification, cells were thawed, resuspended in Lysis Buffer [20 mM HEPES (pH 7.2), 300 mM NaCl, 10% glycerol, 5 mM 2-mercaptoethanol (BME) or 0.5 mM tris(2-carboxyethyl)phosphine (TCEP), 0.25 mM phenylmethylsulfonyl fluoride (PMSF)] (4 °C). The cells were lysed by sonication (4 °C, 75% amplitude, 10 s × 3). The lysate was clarified by centrifugation (15,000 *g*, 30 min, 4 °C). The cleared lysate was mixed with cobalt IMAC resin (Talon resin) and incubated (4 °C, 1 h). The resin was pelleted by centrifugation (700 *g*, 2 min, 4 °C) and washed four times with Wash Buffer [50 mM HEPES (pH 7.2), 300 mM NaCl, 10% glycerol, 5 mM BME, 0.25 mM PMSF, 10 mM imidazole]. His-tagged protein was eluted with Elution Buffer [20 mM HEPES (pH 7.2), 300 mM NaCl, 10% glycerol, 5 mM BME or 0.5 mM TCEP, 0.25 mM PMSF, 100 mM imidazole]. The eluted protein was concentrated with a MW-cutoff spin filter (50 kDa MWCO) and exchanged into Storage Buffer [20 mM HEPES (pH 7.2), 300 mM NaCl, 10% glycerol, 5 mM BME or 0.5 mM TCEP]. The concentration of 8His-MBP-[TEV]-COQ8A^NΔ250^ was determined by its absorbance at 280 nm (ε = 96,720 M^−1^ cm^−1^) (MW = 89.9 kDa) and 8His-MBP-[TEV]-COQ8A^NΔ162^ (ε = 96,720 M^−1^ cm^−1^) (MW = 99.26 kDa). The fusion protein was incubated with Δ238TEV protease (1:50, TEV/fusion protein, mass/mass) (1 h, 20 °C). The TEV protease reaction mixture was mixed with cobalt IMAC resin (Talon resin), incubated (4 °C, 1 h). The resin was pelleted by centrifugation (700 *g*, 3 min, 4 °C). The unbound COQ8A was collected and concentrated with a MW-cutoff spin filter (30 kDa MWCO) and exchanged into storage buffer. The concentration of COQ8A^NΔ250^ was determined by its absorbance at 280 nm (ε = 28,880 M^−1^ cm^−1^) (MW = 45.6 kDa). The concentration of COQ8A^NΔ162^ was determined by its absorbance at 280 nm (ε = 28,880 M^−1^ cm^−1^) (MW = 54.99 kDa). The protein was aliquoted, frozen in N_2(l)_, and stored at –80 °C. Fractions from the protein preparation were analyzed by SDS-PAGE.

*Yeast Coq8p^NΔ41^*. Coq8p^NΔ41^ plasmid constructs were transformed into RIPL competent *E. coli* cells for protein expression. 8His-MBP-[TEV]-Coq8p^NΔ41^ was overexpressed in *E. coli* by autoinduction. Cells were isolated by centrifugation, frozen in N_2(l)_, and stored at −80 °C until further use. For protein purification, cells were thawed on ice, resuspended in Lysis Buffer [50 mM HEPES (pH 7.5), 150 mM NaCl, 5% glycerol, 1 mM BME, 0.25 mM PMSF, 1 mg/mL lysozyme, pH 7.5] and incubated (1 h, 4 °C). The cells were lysed by sonication (4 °C, 6 V, 60 s × 4). The lysate was clarified by centrifugation (15,000 *g*, 30 min, 4 °C). The cleared lysate was mixed with cobalt IMAC resin (Talon resin) and incubated (4 °C, 1 h). The resin was pelleted by centrifugation (700 *g*, 5 min, 4 °C) and washed four times with Wash Buffer [50 mM HEPES (pH 7.5), 150 mM NaCl, 5% glycerol, 1 mM BME, 0.25 mM PMSF, 10 mM imidazole, pH 7.5] (10 resin bed volumes). His-tagged protein was eluted with Elution Buffer [50 mM HEPES (pH 7.5), 150 mM NaCl, 5% glycerol, 1 mM BME, 100 mM imidazole, pH 7.5]. The eluted protein was concentrated with a MW-cutoff spin filter (50 kDa MWCO) and exchanged into storage buffer [50 mM HEPES (pH 7.5), 150 mM NaCl, 5% glycerol, 1 mM BME, pH 7.5]. The concentration of 8His-MBP-[TEV]-Coq8p^NΔ41^ was determined by its absorbance at 280 nm (ε = 109,210 M^−1^ cm^−1^) (MW = 96.2 kDa). The fusion protein was incubated with Δ238TEV protease (1:50, TEV/fusion protein, mass/mass) (1 h, 20 °C). The TEV protease reaction mixture was mixed with cobalt IMAC resin (Talon resin) and incubated (4 °C, 1 h). The unbound Coq8p^NΔ41^ was isolated and concentrated with a MW-cutoff spin filter (30 kDa MWCO) and exchanged into Storage Buffer. The concentration of Coq8p^NΔ41^ was determined by its absorbance at 280 nm (ε = 41,370 M^−1^ cm^−1^) (MW = 52 kDa). The protein was aliquoted, frozen in N_2(l)_, and stored at −80 °C. Fractions from the protein preparation were analyzed by SDS-PAGE.

*Mouse PKA*. 8-His-MBP-PKA and PKA (mouse PKA, Prkaca) were isolated as described above for Coq8p^NΔ41^. The concentration of 8His-MBP-[TEV]-PKA was determined by its absorbance at 280 nm (ε = 121,700 M^−1^ cm^−1^) (MW = 84.8 kDa). The concentration of PKA was determined by its absorbance at 280 nm (ε = 53,860 M^−1^ cm^−1^) (MW = 40.5 kDa).

*E. coli MBP-UbiB^CΔ47^* and MBP. 8His-MBP-[TEV]-UbiB^CΔ47^ and 8His-MBP-[TEV]- were overexpressed in *E. coli* by autoinduction. Cells were isolated and frozen at −80 °C until further use. For protein purification, cells were thawed, resuspended in Lysis Buffer [50 mM HEPES (pH 7.2), 300 mM NaCl, 10% glycerol (w/v), 5 mM BME, 0.25 mM PMSF] (4 °C). The cells were lysed by sonication (4 °C, 75% amplitude, 20 s × 2). The lysate was clarified by centrifugation (15,000 *g*, 30 min, 4 °C). The cleared lysate was mixed with cobalt IMAC resin (Talon resin) and incubated (4 °C, 1 h). The resin was pelleted by centrifugation (700 *g*, 5 min, 4 °C) and washed four times with Wash Buffer [20 mM HEPES (pH 7.2), 300 mM NaCl, 10% glycerol, 5 mM BME or 0.5 mM TCEP, 0.25 mM PMSF, 10 mM imidazole]. His-tagged protein was eluted with Elution Buffer [50 mM HEPES (pH 7.2), 300 mM NaCl, 10% glycerol, 5 mM BME, 0.25 mM PMSF, 100 mM imidazole]. The eluted protein was concentrated with a MW-cutoff spin filter (50 kDa MWCO) and exchanged into storage buffer [50 mM HEPES (pH 7.2), 300 mM NaCl, 10% glycerol, 5 mM BME]. The concentration of 8His-MBP-[TEV]-MBP-UbiB^CΔ47^ was determined by its absorbance at 280 nm (ε = 142,670 M^−1^ cm^−1^) (MW = 102.1 kDa). MBP-[TEV]- (ε = 67,840 M^−1^ cm^−1^) (MW = 44.284 kDa). The protein was aliquoted, frozen in N_2(l)_, and stored at −80 °C. Fractions from the protein preparation were analyzed by SDS-PAGE.

### Nucleotide Binding

The general differential scanning fluorimetry (DSF) method (thermal shift assay) has been described previously (Niesen et al., 2007). Mixtures (20 μL total volume) of Coq8p^NΔ41^ (2 μM) were prepared with SYPRO Orange dye (Life Tech.) (2×), NaCl (150 mM), HEPES (100 mM, pH 7.5), and ligands (e.g. MgATP). Otherwise, the general DSF method, ligand screen, and dissociation constant experiments were conducted as described for COQ8A^NΔ250^ (Stefely et al., 2015). Coq8p^NΔ41^ proteins used for DSF analysis were prepared as described above for COQ8A constructs. SigmaPlot v13.0 was used to calculate *K*_d, app_ and ΔT_m, max_ values using a single site binding model using two replicate experiments.

### ADP-Glo Assay

ADP-Glo (Promega) was performed according to the manufacturer’s instruction with the following modifications. All solutions were diluted in HBS (150 mM NaCl, 20 mM HEPES pH 7.5). Coq8p^NΔ41^ (1 μM), COQ8A^NΔ250^ (2 or 4 μM), COQ8A^NΔ162^ (2 μM), or MBP-UbiB^CΔ47^ (0.5 μM) were mixed with liposomes (~3.33 mM), ATP (100 μM), MgCl_2_ (4 mM), Triton X-100 (1 mM) or reduced Triton X-100 (1 mM) (Sigma Aldrich) and CoQ headgroup analogs (Sigma Aldrich) dissolved as 200 mM stock solutions in DMSO (2-PP, 2-propylphenol; 2-methoxy-6-methylphenol; 4-methylcatechol; 4-HB, 4-hydroxybenzoic acid; 2-MeO-HQ, 2-methoxyhydroquinone; 4-MeOP, 4-methoxyphenol; 2-AP, 2-allylphenol; HQ, hydroquinone; 3,4-di-HB, 3,4-dihydroxybenzoic acid; MeHQ, methylhydroquinone; 2,6-diMeO-HQ, 2,6-dimethoxyhydroquinone) (1 mM) (final concentrations for reaction components). Reaction were incubated (30 °C, 45 min). Then ADP-Glo Reagent (5 μL) was added and incubated [40 min, r.t. (~21 °C), covered]. Kinase Detection Reagent was added (10 μL) and incubated (40 min, r.t., covered). Luminescence was read using default values. An ADP/ATP standard curve was made according to the manufacturer’s instructions. Error bars represent s.d. of technical triplicate measurements unless otherwise specified.

### Cytophos ATPase Assay (Malachite Green)

The Cytophos ATPase assay was performed according to the manufacturer’s instruction with the following modifications. All solutions were diluted in HBS (150 mM NaCl, 20 mM HEPES pH 7.5). COQ8A^NΔ250^ (1 μM final), Coq8p^NΔ41^ (1 μM final), COQ8A^NΔ162^ (1 μM final) or MBP-UbiB^CΔ47^ (0.25 μM final) with 100 μM ATP, 4 mM MgCl_2_, 1 mM 2-alkylphenol, 0.5 mM CoQ_1_, 1 mM Triton X-100 or reduced Triton X-100. CoQ_1_ was reduced by adding 1.2 fold molar excess of NaBH_4_ (Sigma) (10 min, r.t.). Reactions were incubated (30 °C, 45 min) and 35 μL of Cytophos reagent was added and incubated (10 min, r.t.). Absorbance was measured at 650 nm. Reactions in Figure 2E were incubated for 10 min rather than 45 min and [Coq8p^NΔ41^] was 0.5 μM. The standard curve ranged from 0–50 μM phosphate. Error bars represent s.d. of technical triplicate measurements unless otherwise specified.

### *In Vitro* [γ-^32^P]ATP ATPase Assay

Method for Figures S4D and S4E.

Unless otherwise indicated, COQ8A^NΔ162^, Coq8p^NΔ41^ (4 μM), or PKA (0.2 μM) was mixed with [γ-^32^P]ATP (0.25 μCi/μL, 100 μM [ATP]_total_) and MgCl_2_ (4 mM) in an aqueous buffer (20 mM HEPES, 150 mM NaCl, pH 7.5) and incubated (30 °C, 45 min, 700 rpm) (final concentrations for reaction components). Myelin basic protein (1 μM) and a cell free expressed COQ protein complex consisting of COQ3, COQ4, COQ5, COQ6, COQ7-strep, and COQ9 (~1.5–0.1 μM). Half (10 μL) of each reaction was quenched with 4× LDS buffer. [γ-^32^P]ATP was separated from COQ8 by SDS-PAGE (10% Bis-Tris gel, MES buffer, 150 V, 1.5 h). The gel was stained with Coomassie Brilliant Blue, dried under vacuum, and imaged by digital photography. The other half (10 μL) of the reaction was quenched with 1:1 CHCl_3_:MeOH (50 μL, 4 °C) and 1 M HCl (12.5 μL, 4 °C). Reactions were mixed by vortexing (3 × 10 s), centrifuged (3000 *g*, 1 min, 4 °C) and the aqueous layer was discarded. 10 μL of the organic layer was spotted on silica TLC and lipids were separated with CHCl_3_/MeOH/30% NH_4_OH_(aq)_/H_2_O (50:40:8:2, v/v/v/v). A storage phosphor screen was exposed to the gel or TLC plate (~5 days) and then imaged with a Typhoon (GE) to generate the phosphorimages.

### *In Vitro* [γ-^32^P]ATP ATPase Assay

Unless otherwise indicated, COQ8A^NΔ162^ (2 μM) or Coq8p^NΔ41^ (1 μM) was mixed with liposomes (~3.33 mM), [γ-^32^P]ATP (0.01 μCi/μL, 100 μM [ATP]_total_), and MgCl_2_ (4 mM) in an aqueous buffer (20 mM HEPES, 150 mM NaCl, pH 7.5) and incubated (20 μL total volume, 30 °C, 45 min, 700 rpm) (final concentrations for reaction components). Reactions were quenched with 0.75 M potassium phosphate pH 3.3 (20 μL, 4 °C). 1 μL of quenched reaction was spotted on PEIcellulose TLC (Millipore) and developed with 0.5 M LiCl in 1 M formic acid_(aq)_. After drying, a storage phosphor screen was exposed to the PEI-cellulose TLC plate (~5 hours) and then imaged with a Typhoon (GE) to generate the phosphorimages.

### Yeast Drop Assay and Growth Curves

Yeast cultures for drop assays. *S. cerevisiae* (W303) Δ*coq8* yeast were transformed as previously described (Gietz and Woods, 2002) with p426 GPD plasmids encoding Coq8p variants and grown on uracil drop-out (Ura^−^) synthetic media plates containing glucose (2%, w/v). Individual colonies of yeast were used to inoculate Ura^−^ media (2%, w/v) starter cultures, which were incubated (30 °C, ~18 h, 230 rpm). Serial dilutions of yeast (10^4^, 10^3^, 10^2^, or 10 yeast cells) were dropped onto Ura^−^ agar media plates containing either glucose (2%, w/v) or glycerol (3%, w/v) and incubated (30 °C, 2 d). To assay yeast growth in liquid media, yeast from a starter culture were swapped into Ura^−^ media with glucose (0.1%, w/v) and glycerol (3%, v/v) at an initial density of 5×10^6^ cells/mL. The cultures were incubated in a sterile 96 well plate with an optical, breathable cover seal (shaking at 1140 rpm). Optical density readings were obtained every 10 min. For Coq8p-*AS1* experiments, individual colonies of *S. cerevisiae* (BY4742) were used to inoculate Ura^−^ media (20 g/L glucose) starter cultures, which were incubated (30 °C, ~12 h, 230 rpm). Yeast were diluted to 2.5×10^4^ cells/mL and incubated until early log phase (30 °C, 16 h, 230 rpm). Yeast were swapped into Ura^−^ media with glucose (0.1%, w/v) and glycerol (3%, v/v) at an initial density of 5×10^6^ cells/mL with or without 50 μM 3-MB-PP1. The cultures were incubated in a sterile 96 well plate with an optical, breathable cover seal (shaking at 1140 rpm). Optical density readings were obtained every 10 min.

### Growth of Yeast and Extraction of Yeast CoQ_6_ for LC-MS Quantitation

Method for Figure 2D adapted from (Stefely et al., 2015). 2.5×10^6^ yeast cells (as determined by OD_600_ of a starter culture) were used to inoculate a 25 mL culture of Ura^−^ media (10 g/L glucose), which was incubated (30 °C, 230 rpm) for 23 h. At 23 h, the yeast cultures were ~4 h past the diauxic shift and the media was depleted of glucose. The OD_600_ of the culture was measured and used to determine the volume of culture needed to isolate 1×10^8^ yeast cells. 1×10^8^ yeast cells were pelleted by centrifugation in 15 mL conical tubes (4,000 *g*, 3 min, 4 °C) and transferred to 1.5 mL tubes and spun again (16,000 *g*, 0.5 min, 4 °C), the supernatant was discarded, and the yeast pellet was flash frozen in liquid N_2_ and stored at −80 °C. A frozen pellet of yeast (10^8^ yeast cells) was thawed on ice and mixed with phosphate buffered saline (200 μL) and glass beads (0.5 mm diameter, 100 μL). The yeast were lysed by vortexing with the glass beads (30 s). Coenzyme Q_10_ (CoQ_10_) was added as an internal standard (10 μM, 10 μL), and the lysate was vortexed (30 s). Hexanes/2-propanol (10:1, v/v) (500 μL) was added and vortexed (2 × 30 s). The samples were centrifuged (3,000 *g*, 1 min, 4 °C) to complete phase separation. 400 μL of the organic phase was transferred to a clean tube and dried under Ar_(g)_. The organic residue was reconstituted in ACN/IPA/H_2_O (65:30:5, v/v/v) (100 μL) by vortexing (30 s) and transferred to a glass vial for LCMS analysis. Samples were stored at −80 °C.

For analog-sensitive *coq8* experiments (Figure 6D), 2.75×10^6^ yeast cells (as determined by OD_600_ of a starter culture) were used to inoculate a 75 mL culture of Ura^−^, *p*ABA^−^ media (20 g/L glucose), which was incubated (30 °C, 230 rpm) for 16 h. Yeast were diluted to 5×10^6^ cells/mL (as determined by OD_600_ of a starter culture) in respiratory Ura^−^, *p*ABA^−^ media (0.1 % glucose, w/v; 3% glycerol, w/v) and incubated (30 °C, 230 rpm) for 24 h in the presence and absence of 25 μM 3-MB-PP1 inhibitor. The OD_600_ of the culture was measured and used to determine the volume of culture needed to isolate 5×10^7^ yeast cells. 5×10^7^ yeast cells were pelleted by centrifugation in 15 mL conical tubes (4,000 *g*, 3 min, 4 °C) and transferred to 1.5 mL tubes and spun again (16,000 *g*, 0.5 min, 4 °C), the supernatant was discarded, and the yeast pellet was flash frozen in liquid N_2_ and stored at −80 °C. A frozen pellet of yeast (5×10^7^ yeast cells) was thawed on ice and mixed with phosphate buffered saline (200 μL) and glass beads (0.5 mm diameter, 100 μL). The yeast were lysed by vortexing with the glass beads (30 s). Coenzyme Q_10_ (CoQ_10_) was added as an internal standard (10 μM, 10 μL), and the lysate was vortexed (30 s). Cold CHCl_3_/MeOH (1:1, v/v) (900 μL) was added and vortexed (2 × 30 s). Samples were acidified with HCl (1 M, 200 μL) and vortexed (2 × 30 s). The samples were centrifuged (5,000 *g*, 2 min, 4 °C) to complete phase separation. 400 μL of the organic phase was transferred to a clean tube and dried under Ar_(g)_. The organic residue was reconstituted in ACN/IPA/H_2_O (65:30:5, v/v/v) (100 μL) by vortexing (2 × 30 s) and transferred to a glass vial for LC-MS analysis. Samples were stored at −80 °C.

### Targeted LC-MS for Yeast CoQ_6_

Method for Figure 2D. LC-MS analysis was performed on an Acquity CSH C18 column held at 50 °C (100 mm × 2.1 mm × 1.7 μL particle size; Waters) using (400 μL/min flow rate; Thermo Scientific). Mobile phase A consisted of 10 mM ammonium acetate in ACN/H_2_O (70:30, v/v) containing 250 μL/L acetic acid. Mobile phase B consisted of 10 mM ammonium acetate in IPA/ACN (90:10, v/v) with the same additives. Initially, mobile phase B was held at 40% for 6 min and then increased to 60% over 3 min followed by an increase to 85% over 15 s. Mobile phase B was then increased to 99% over 75 s where it was held for 30 s. The column was re-equilibrated for 4 min before the next injection. 10 μL of sample was injected for each sample. The LC system was coupled to a Q Exactive mass spectrometer (Build 2.3 SP2) by a HESI II heated ESI source kept at 300 °C. The inlet capillary was kept at 300 °C, sheath gas was set to 25 units, and auxiliary gas to 10 units. For identification of CoQ_6_ and CoQ_10_ species, the MS was operated in positive mode (5 kV) and masses (591.44 and 880.71, respectively) were targeted for fragmentation. AGC target was set to 1×10^6^ and resolving power was set to 140,000. Quantitation was performed in Xcalibur (Thermo) by monitoring the product ion 197.08 Th, corresponding to the Q headgroup, for each targeted mass.

For Coq8p-*AS1* lipid samples (Figure 6D), LC-MS analysis was performed on an Acquity CSH C18 column held at 50 °C (100 mm × 2.1 mm × 1.7 μL particle size; Waters) using an Ultimate 3000 RSLC Binary Pump (400 μL/min flow rate; Thermo Scientific). Mobile phase A consisted of 10 mM ammonium acetate in ACN/H_2_O (70:30, v/v) containing 250 μL/L acetic acid. Mobile phase B consisted of 10 mM ammonium acetate in IPA/ACN (90:10, v/v) with the same additives. Mobile phase B was held at 50% for 1.5 min and then increased to 95% over 6.5 min where it was held for 2 min. The column was then reequilibrated for 3.5 min before the next injection. 10 μL of sample were injected by an Ultimate 3000 RSLC autosampler (Thermo Scientific). The LC system was coupled to a Q Exactive mass spectrometer by a HESI II heated ESI source kept at 325 °C (Thermo Scientific). The inlet capillary was kept at 350 °C, sheath gas was set to 25 units, auxiliary gas to 10 units, and the spray voltage was set to 3,000 V. The MS was operated in negative polarity mode acquiring MS^1^ spectra from 200–1550 m/z supplemented with scheduled targeted scan modes to quantify key CoQ intermediates. The resulting LC-MS data were processed using TraceFinder 4.0 (Thermo Fisher Scientific). Metabolite signals were integrated and normalized to the CoQ_10_ internal standard. Error bars represent s.d. across three biological replicates.

### AP-MS Lipidomics of 8His-MBP-[TEV]-MBP-UbiB^CΔ47^

8His-MBP-[TEV]-MBP-UbiB^CΔ47^ protein purifications were performed in biological triplicate according to the method used above. Eq/S buffer [50 mM HEPES (pH 7.2), 300 mM NaCl, 10% glycerol (w/v)] was added to 25 nmol of 8His-MBP-[TEV]-MBP-UbiB^CΔ47^ to bring total sample volume to 682 μL. 20 μL of 25 μM CoQ_6_ internal standard (in CHCl_3_:MeOH, 1:1, v/v) (Avanti Polar Lipids) was added to each sample and vortex 10 s (500 pmols) (protein precipitated upon mixing). Cold CH_3_Cl:MeOH (1:1, v/v, 4.3 mL) was added to each sample and mixed to form one phase by vortexing (1 × 30 s, 4 °C). Samples were acidified by adding HCl (1 M, 30 μL) and vortexing (2 × 30 s). Phase separation was induced by addition of saturated NaCl_(aq)_ (100 μL) and mixed by vortexing (10 s). Phase separation was completed by centrifugation (3000 *g*, 1 min, 4 °C). Aqueous layer was discarded and 2.9 mL of the organic layer was transferred to new glass tube and dry under Ar_(g)_ (r.t., ~2 hr). The organic residue was reconstituted in ACN/IPA/ H_2_O (65:30:5, v/v/v) (200 μL) by vortexing (30 s, 4 °C), bath sonication (60 s, 4 °C), and vortexing (30 s, 4 °C). Transfer 180 μL to an autosampler vial labeled, stored under Ar_(g)_ at −80 °C.

### LC-MS Discovery and Targeted Methods for AP-MS

Both targeted and discovery-based lipidomics of UbiB were done using the instrumentation, column, and mobile phases described in Targeted LC-MS for yeast CoQ_6_.

*Discovery Lipidomics*. Mobile phase B started at 2% and increased to 85% over 20 min, then increased to 99% over 1 min, where it was held for 7 min. The column was then reequilibrated for 2 min before the next injection. The LC system was coupled to a Q Exactive mass spectrometer by a HESI II heated ESI source kept at 300 °C (Thermo Scientific). The inlet capillary was kept at 300 °C, sheath gas was set to 25 units, auxiliary gas to 10 units, and the spray voltage was set to 3,000 V. The MS was operated in polarity switching mode, acquiring both positive mode and negative mode MS^1^ and MS^2^ spectra. The AGC target was set to 1×10^6^ and resolving power to 17,500. Ions from 200-1600 m/z were isolated (Top 2) and fragmented by stepped HCD collision energy (20, 30, 40). The resulting LC–MS/MS data were processed using Compound Discoverer 2.0 (Thermo Fisher) and an in-house-developed software suite. Briefly, all m/z peaks were aggregated into distinct chromatographic profiles (i.e., compound groups) using a 10 p.p.m. mass tolerance. These chromatographic profiles were then aligned across all LC-MS/MS experiments using a 0.2 min retention time tolerance. All compound groups were compared against a matrix run and only compound groups whose intensity was 5-fold greater were retained. MS/MS spectra were searched against an in-silico generated lipid spectral library containing 35,000 unique molecular compositions representing 31 distinct lipid classes. Spectral matches with a dot product score greater than 650 and a reverse dot product score greater than 750 were retained for further analysis. Identifications were further filtered using precursor mass, MS/MS spectral purity, retention time, and chromatographic profile. Lipid MS/MS spectra which contained no significant interference (<75%) from co-eluting isobaric lipids were identified at the individual fatty acid substituent level of structural resolution. Otherwise, lipid identifications were made with the sum of the fatty acid substituents. All lipid intensities were normalized to the total intensity of all lipids in the chromatogram following the void volume. Intensities of each individual lipid were divided by the sum of the intensities for each sample and averaged across all three biological replicates. Student’s *t*-test was used to determine statistical significance. Nomenclature for lipid acyl chains is from (Liebisch et al., 2013).

*Targeted Lipidomics for CoQ and its Intermediates*. LC-MS analysis was performed on an Acquity CSH C18 column held at 50 °C (100 mm × 2.1 mm × 1.7 μL particle size; Waters) using an Ultimate 3000 RSLC Binary Pump (400 μL/min flow rate; Thermo Scientific). Mobile phase A consisted of 10 mM ammonium acetate in ACN/H_2_O (70:30, v/v) containing 250 μL/L acetic acid. Mobile phase B consisted of 10 mM ammonium acetate in IPA/ACN (90:10, v/v) with the same additives. Mobile phase B was held at 50% for 1.5 min and then increased to 95% over 6.5 min where it was held for 2 min. The column was then reequilibrated for 3.5 min before the next injection. 10 μL of sample were injected by an Ultimate 3000 RSLC autosampler (Thermo Scientific). The LC system was coupled to a Q Exactive mass spectrometer by a HESI II heated ESI source kept at 300 °C (Thermo Scientific). The inlet capillary was kept at 300 °C, sheath gas was set to 25 units, auxiliary gas to 10 units, and the spray voltage was set to 4,000 V and 5,000 V for positive and negative mode respectively. CoQ and its intermediates were targeted for quantification. The MS was operated in positive or negative mode depending on the intermediate being targeted. Metabolite signals were integrated and normalized to a CoQ_6_ internal standard. Error bars represent s.d. of biological triplicate measurements.

Table of targeted species for AP-MS experiments.

| Species | Mass (m/z) | Polarity | Start (min) | End (min) | (N)CE | (N)CE type |
| --- | --- | --- | --- | --- | --- | --- |
| CoQ<sub>8</sub> | 727.566 | Positive | 5.5 | 6 | 27 | NCE |
| CoQ<sub>6</sub> | 591.44 | Positive | 3.6 | 4.1 | 27 | NCE |
| OPP | 637.5354 | Negative | 5.7 | 6.15 | 30 | NCE |
| OHB | 681.5398 | Negative | 4.3 | 5 | 30 | NCE |

### Liposomes

Liposomes were made by drying down lipids in a 5 mL plastic tube under Ar_(g)_ at room temperature until a film was left. Liposomes were made of the following lipid (Avanti Polar Lipids) compositions: PC, PC/NBD-PE 99.9/0.1; PC/PE, PC/PE/NBD-PE 89.9/10/0.1; PC/CL, PC/CL/NBD-PE 89.9/10/0.1; PC/PE/CL, PC/PE/CL/NBD-PE 79.9/10/10/0.1; PC/CoQ_10_, PC/CoQ_10_/NBD-PE 97.9/2/0.1; PC/PG, PC/PG/NBD-PE 89.9/10/0.1; PC/PS, PC/PS/NBD-PE 89.9/10/0.1; PC/PA, PC/PA/NBD-PE 89.9/10/0.1; PC/CDP-DAG, PC/CDP-DAG/NBD-PE 89.9/10/0.1; all mol %. The lipids were dried in a vacuum chamber overnight at 25-30 inHg vacuum. The dry lipid film was reconstituted in HBS buffer (20 mM HEPES pH 7.5, 150 mM NaCl) at 30–35 °C for one hour with occasional pipetting to resuspend the lipid film. The total concentration of lipids in solution was 10 mM. 2 μL of the liposomes were taken before and after extrusion and diluted with 22 μL of HBS to determine how much the liposomes were diluted during extrusion by measuring the fluorescence of NBD-PE (excitation: 460 nm, emission: 535 nm). Liposomes were extruded through 100 nM membranes (Avanti Polar Lipids), 15 passes, 30–35 °C. Liposomes were made fresh before each experiment.

### Liposome Flotation Assay

Liposome flotation assay is adapted from (Connerth et al., 2012) with the following modifications. Liposomes (100 μL) were mixed with protein (50 μL 6× COQ8) at 4 °C then incubated (r.t., ~10 min). Final concentrations of reagents are as follows: 2–4 μM protein and 3.33 mM liposomes in HBS (150 mM NaCl, 20 mM HEPES pH 7.5). 2.72 M sucrose (110 μL) was added to the protein liposome mixture (1.15 M [sucrose] final). The sucrose-liposome-protein mixture (250 μL) was added to the ultracentrifuge tube. The sucrose gradient was made by layering 300 μL HBS 0.86 M sucrose, 250 μL HBS 0.29 M sucrose, and 150 μL HBS on top of the sucrose-liposome-protein mix. Centrifuge (240,000 *g*, 1 h, 4 °C). Remove 450 μL for the top layer and 450 μL for the bottom layer. Determine liposome location by mixing 8 μL of top or bottom fraction with 16 μL HBS and reading NBD-PE fluorescence for (excitation: 460 nm, emission: 535 nm). To concentrate the protein from the top and bottom fractions, a CHCl_3_:MeOH precipitation was performed according to (Wessel and Flugge, 1984). Add 1800 μL methanol (4 volumes) to a 450 μL fraction. After thorough mixing 450 μL chloroform (one volume) is added. Vortex then add 1350 μL water (three volumes), vortex again and centrifuge immediately for 5 minutes at full speed in a microfuge. A white disc of protein should form between the organic layer (bottom) and the aqueous layer (upper). Discard most of the upper aqueous layer and be careful not to disturb the protein pellet. Add 1000 μL of methanol to the tube and invert 3 times. Centrifuge for 5 minutes at full speed (16,000 *g*). Remove all liquid and allow the pellet to air dry. The precipitated protein pellet was resuspend in 30 μL 1× LDS with 10 mM DTT, incubated at 95 °C ~10 min, and analyzed with 4–12% Novex NuPAGE Bis-Tris SDS-PAGE (Invitrogen) gels (1 hr, 150 V). Band quantification was done with imaging and densitometry on a LiCOR Odyessey CLx (700 nm) using Image Studio v5.2 software. Error bars for Figure 4G represent s.d. of three separate floats. Student’s *t*-test was used to determine statistical significance.

### QUANTIFICATION AND STATISTICAL ANALYSIS

See each individual method for the associated statistical analysis.

## SUPPLEMENTAL MOVIES AND TABLES

**Movie S1, related to Figure 4. CG-MD simulation of COQ8A with PC bilayer.**

4 microseconds of production run of a CG-MD simulation of COQ8A (green) (PDB:4PED) with PC bilayer (phosphate heads in gray). The positively charged residues R262, R265, K269 are colored blue.

**Movie S2, related to Figure 4. CG-MD simulation of COQ8A with PC/PE/CL bilayer.**

4 microseconds of production run of a CG-MD simulation of COQ8A (green) (PDB:4PED) with PC/PE/CL bilayer (PC/PE phosphate heads in gray and CL phosphate heads in red). The positively charged residues R262, R265, K269 are colored blue.

**Table S1. Explanation of UbiB family member variants used in this study.**

**Table S2, related to Figure 1. UbiB AP-MS lipidomics data.**

**Table S3, related to Figure 1. List of compounds and hits from NMR line-broadening screen.**

**Table S4. Oligonucleotide primers used in this study.**

## SUPPLEMENTAL INFORMATION

**Figure S1, related to Figure 1.**
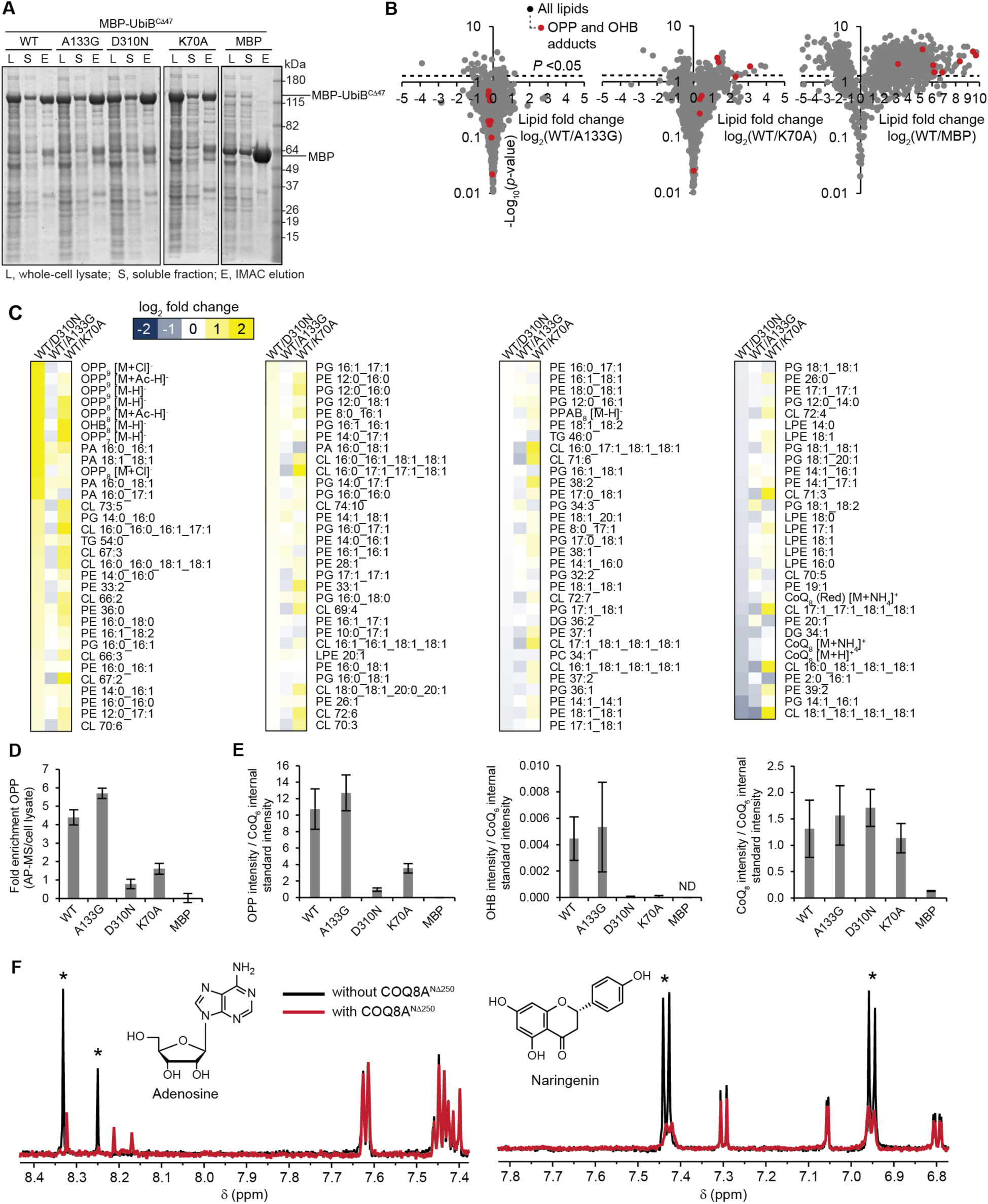
UbiB family members bind CoQ precursor-like lipids and small molecules. (A) Representative Coomassie stained SDS-PAGE gel from MBP-UbiB^CΔ47^ protein purification for lipidomics and activity assays. (B) Fold changes in lipid abundances (log_2_[(WT MBP-UbiB^CΔ47^)/(mutant or MBP)], n = 3) versus statistical significance as quantified by LC-MS/MS. (C) Heat map of the fold changes for all identified and quantified lipids comparing WT MBP-UbiB^CΔ47^ to mutants. (D) Fold enrichment of OPP with purified protein compared to OPP abundance in matched cell lysate samples. (E) Amount of copurifying OPP, OHB, and CoQ_8_, measured using targeted LC-MS/MS. Error bars in D and E represent s.d. of biological replicate measurements. (F) A selected region of the ^1^H-NMR spectra for mixtures containing adenosine (left) and naringenin (right) with and without COQ8A^NΔ250^. Asterisks (*) indicate significant peaks for ADP or naringenin that exhibit line-broadening. Other peaks belong to different compounds in the mixtures that are not interacting with COQ8A^NΔ250^.

**Figure S2, related to Figure 2.**
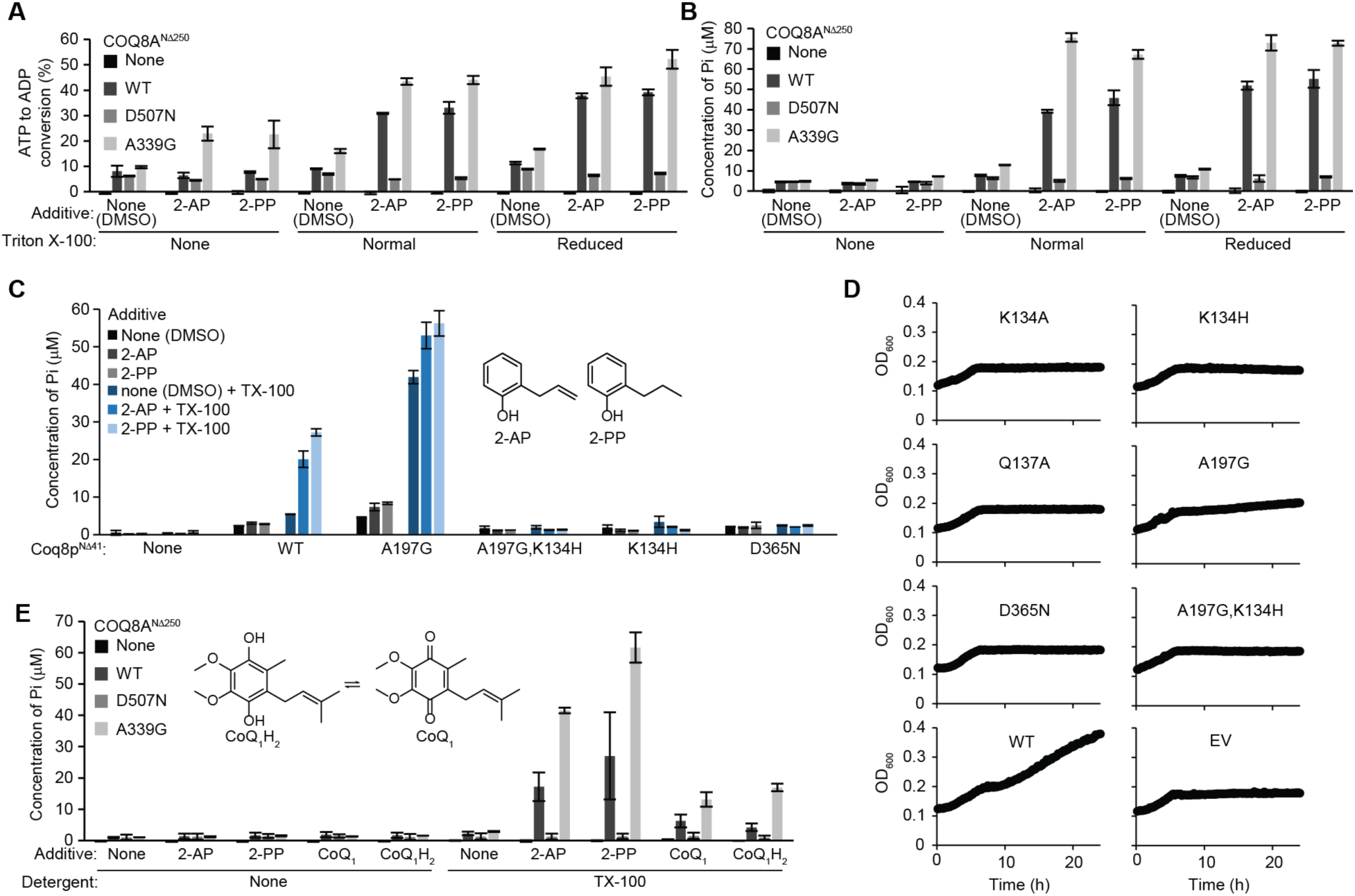
UbiB family members are activated by Triton X-100 and 2-alkylphenols. (A) ADP-Glo assay with COQ8A^NΔ250^ variants, Triton X-100 (TX-100) or Triton X-100 reduced, and 2- alkylphenols.(B) Malachite green assay with COQ8A^NΔ250^ variants, Triton X-100 or Triton X-100 reduced, and 2-alkylphenols. Pi, inorganic phosphate. (C) Malachite green ATPase assay with Coq8p KxGQ mutants, 2-alkylphenols and Triton X-100. (D) Growth curves of Δ*coq8* transformed with KxGQ mutants in respiratory media (0.1% glucose 3% glycerol). (E) Malachite green ATPase assay with COQ8A 2-alkylphenols and CoQ_1_.

**Figure S3, related to Figure 3.**
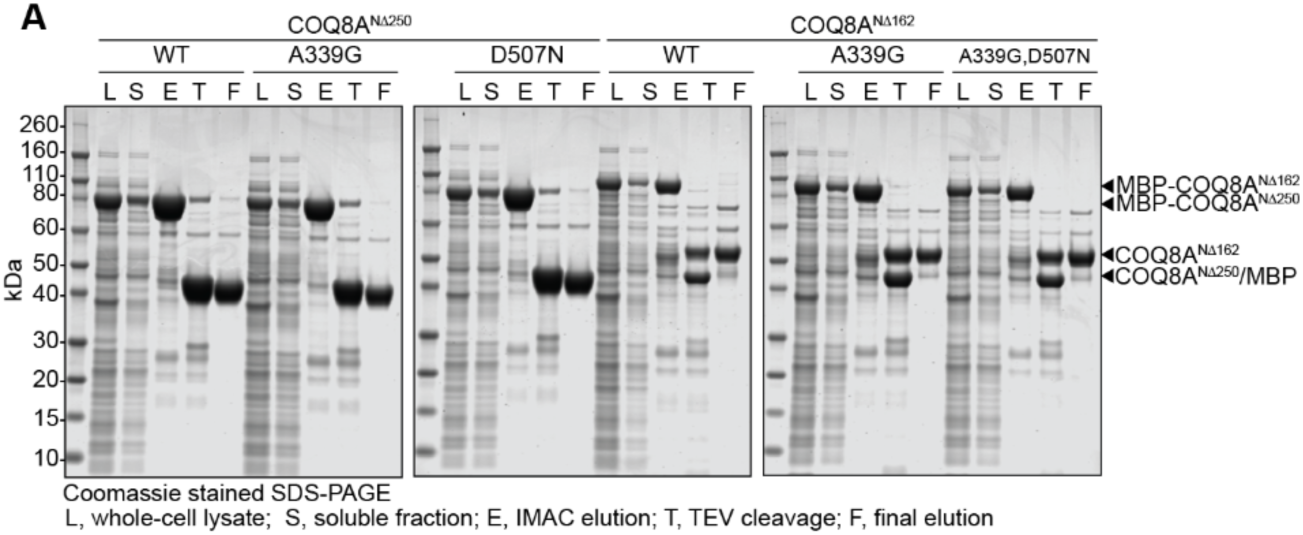
Purification of COQ8A^NΔ162^. (A) Coomassie stained SDS-PAGE of samples from a COQ8A^NΔ250^ and COQ8A^NΔ162^ protein purification.

**Figure S4, related to Figure 4.**
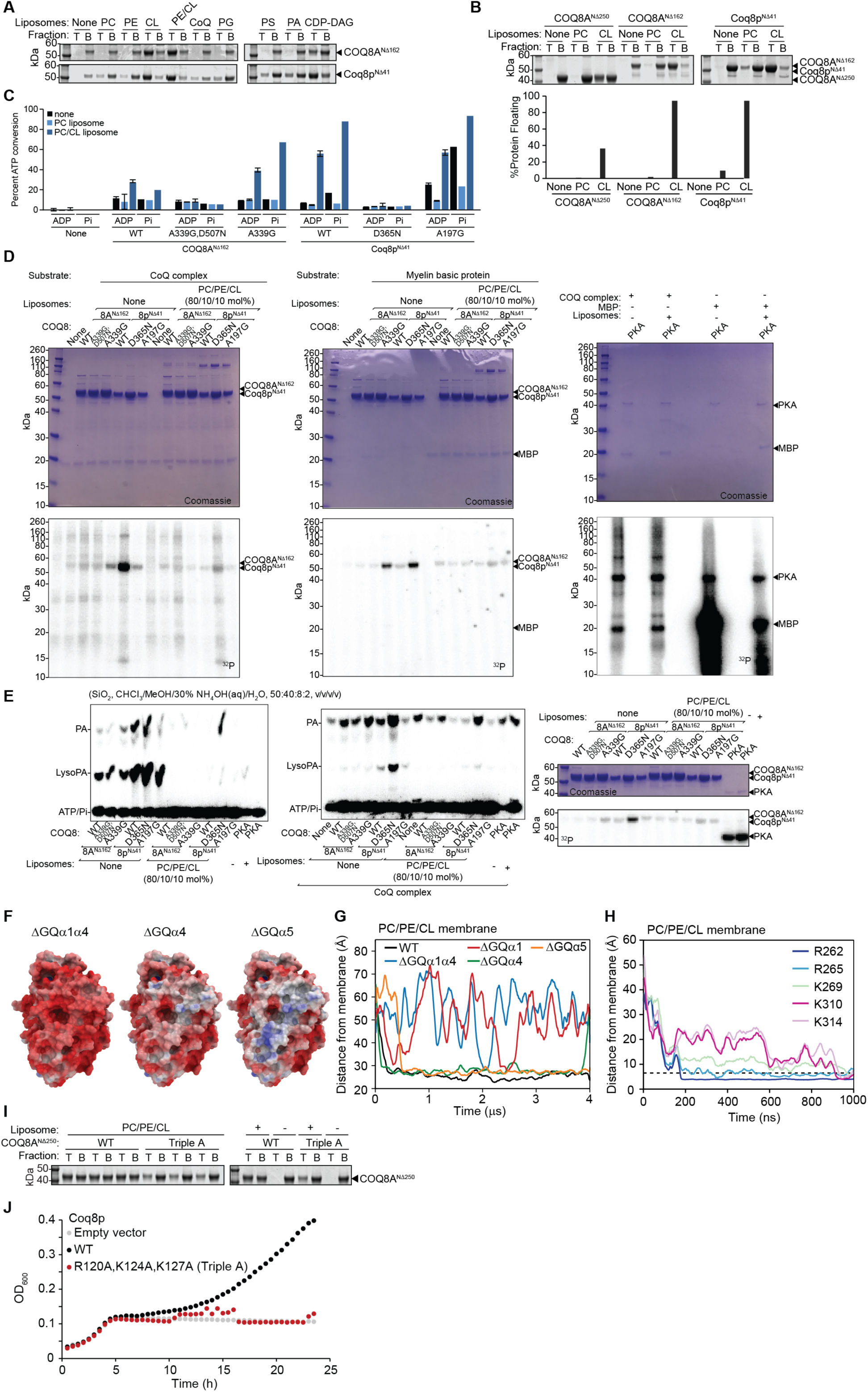
CL enhances the ATPase activity of COQ8 and liposome binding. (A) Coomassie stained SDS-PAGE of the liposome flotation assay in Figure 4A with mature COQ8 and a panel of liposomes. T, top fraction; B, bottom fraction. (B) Coomassie stained SDS-PAGE of the liposome flotation assay with COQ variants and PC/PE/CL liposomes. T, top fraction; B, bottom fraction. (C) ADP-Glo (ADP produced) and [γ-^32^P]ATP ATPase assay (Pi produced) of COQ8 variants with PC or PC/CL liposomes performed in parallel. The amount of Pi produced was calculated from the TLC plates shown beneath the bar graphs. (D) Coomassie stained SDS-PAGE and corresponding phosphorimage of [γ-^32^P]ATP kinase reactions. The Coq8p D365N mutant looks active because it was overloaded. (E) Phosphorimage of silica TLC from [γ-^32^P]ATP kinase reactions. (F) Electrostatic maps of COQ8A with the positively charged or polar residues on helices GQα1, GQα4, and GQα5 mutated to alanine [negative (−5 kcal/e.u. charge): red, via white, to positive (+5 kcal/e.u. charge): blue]. (G) Time evolution of the distance between the center of mass of the protein and the center of mass of the phosphate heads of the leaflet it interacts with, for CG-MD simulations of different GQα mutated COQ8A proteins. (H) Time evolution of the distance between positively charged residues on COQ8A GQα1 and GQα4 and the closest phosphate head. (I) Coomassie stained SDS-PAGE of top and bottom fraction from the liposome flotation assay in Figure 4G. +, with PC/PE/CL liposomes; -, no liposomes. (J) Growth curve of Δ*coq8* transformed with WT *coq8* or the Triple A mutant in respiratory media (0.1% glucose 3% glycerol).

**Figure S5, related to Figure 5.**
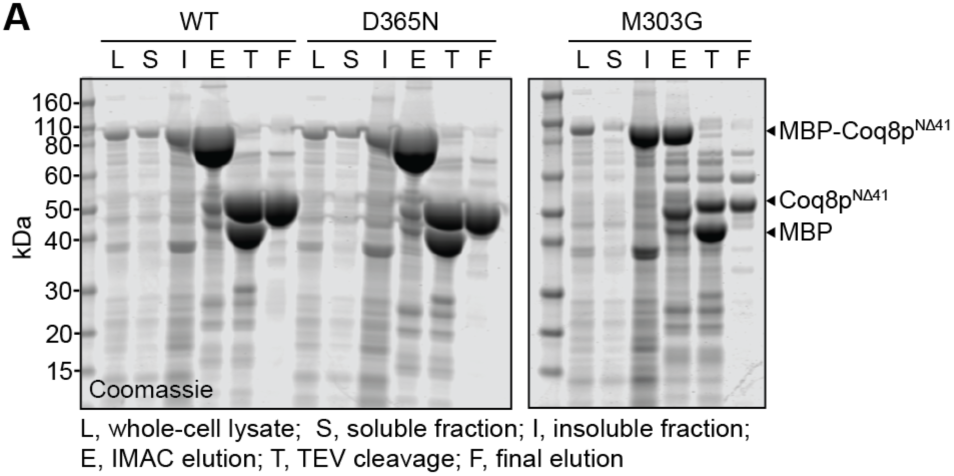
Purification of *as*-Coq8p^NΔ41^. (A) Coomassie stained SDS-PAGE of samples from *as-*Coq8p^NΔ41^ protein purification.

**Figure S6, related to Figure 6.**
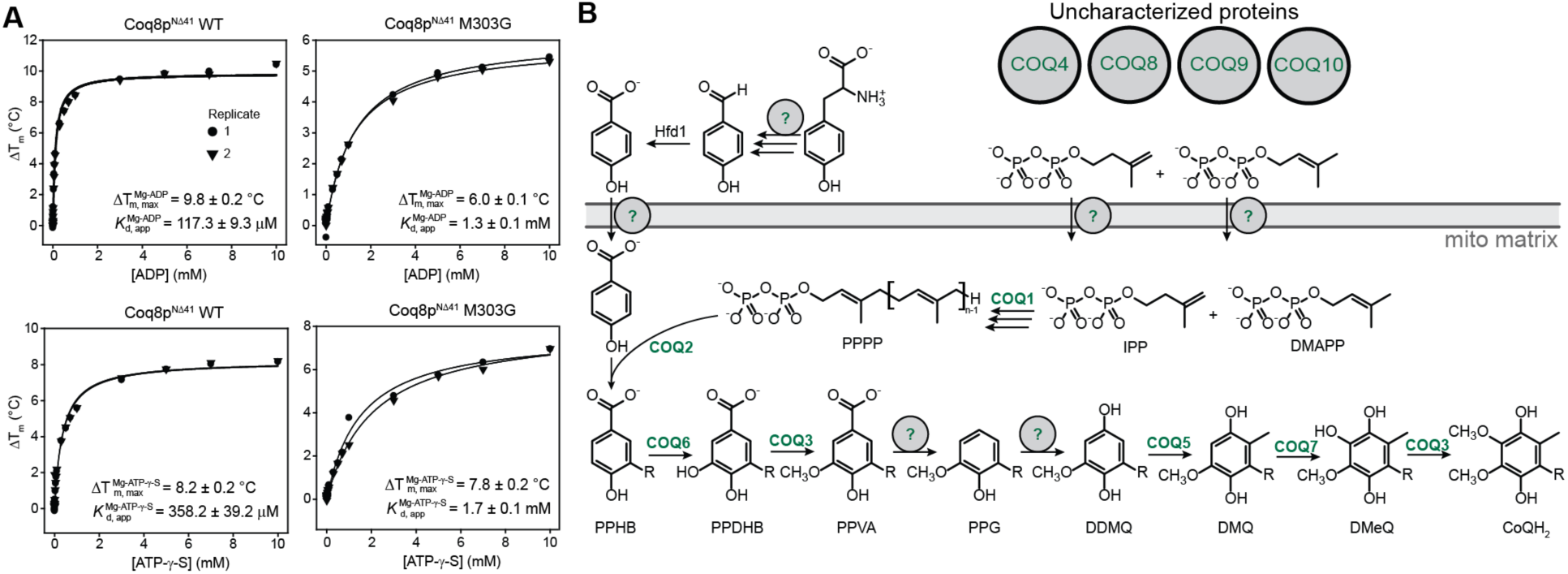
Nucleotide binding analysis of *as-*Coq8p^NΔ41^. (A) MgADP and MgATP-γ-S binding curves for WT and *as*-Coq8p^NΔ41^. (B) Eukaryotic CoQ biosynthesis pathway.

